# Zero-shot virtual knock-out rankings do not recover causal genes from observational single-cell data

**DOI:** 10.64898/2026.11.21.768293

**Authors:** Jiale Cui, Meixuan Li, Rui Zhou, Haoyu Zhang, Saihang Xu, Lei Li, Rong Zhang

**Affiliations:** Department of Obstetrics and Gynecology, Shanxi Provincial People’s Hospital Affiliated to Shanxi Medical University, Taiyuan 030012, Shanxi, China; The Fifth Clinical Medical College of Shanxi Medical University, Taiyuan 030012, Shanxi, China; Department of Radiation Oncology, Shanxi Provincial People’s Hospital Affiliated to Shanxi Medical University, Taiyuan 030012, Shanxi, China; School of Nursing, Shanxi Medical University, Taiyuan 030012, Shanxi, China; National Regional Children’s Medical Center (Northwest); Key Laboratory of Precision Medicine to Pediatric Diseases of Shaanxi Province; Xi’an Key Laboratory of Children’s Health and Diseases, Shaanxi Institute for Pediatric Diseases; Xi’an Children’s Hospital, Affiliated Children’s Hospital of Xi’an Jiaotong University, Xi’an 710003, Shaanxi, China

**Keywords:** virtual knock-out, gene regulatory network, single-cell transcriptomics, causal inference, target prioritisation, DepMap essentiality, network structure, method benchmarking

## Abstract

**Background:** Virtual knock-out methods rank genes by the change predicted after deleting one gene in silico and nominate targets without perturbation data. That use assumes the ranking reflects the causal consequence of removal; we tested it for three zero-shot methods in four configurations under one criterion.

**Results:** We compared scTenifoldKnk (R and Python), CellOracle and Geneformer on 438 real, 3,585 synthetic and 330 DepMap knock-outs. Top-20 lists of different knock-outs over-lapped by a median of 12 to 14 of 20, against 0.08 by chance. Displacement followed out-degree (*R*^2^ = 0.825; 0.852 with six further features). Decoupling causal effect from topology left criterion value and grade unchanged (36 of 36 RED). For scTenifoldKnk, the ranking showed no evidence of recovery above chance where causal genes were planted (AUROC 0.538 and 0.528 in the two constructions). CellOracle’s displacement showed no detectable relation to the data-built network (*ρ* = *−*0.212, *P* = 0.556). Geneformer’s shift tracked token coverage inconsistently (+0.41, +0.40, *−*0.27). Against DepMap essentiality, scTenifoldKnk’s Python implementation (scTenifoldpy 0.5.1) reached AUROC 0.459 (95% CI 0.357–0.566), not distinguishable from chance; detection rate reached 0.588 (95% CI 0.497–0.677).

**Conclusions:** For scTenifoldKnk and CellOracle, the ranking reports network properties, not the causal consequence of removal. No measured structural feature accounted for more than a small fraction of Geneformer’s shift. Detection rate, needing no network, gained +0.129 (95% CI 0.0017–0.2543; 97.6% of resamples positive) over the tool.

## 1 Background

Removing one gene and asking what changes downstream is a routine question in single-cell studies, and it is usually answered without an experiment. A virtual knock-out is any method that predicts the downstream consequence of removing or reducing one gene’s expression in a model fitted to observational data. The output is used to rank genes for follow-up work: to prioritise targets within a cell type, to interpret a pathway, and to nominate candidates for drug repurposing. Studies of lineage conversion [1], of hepatocyte identity in liver cancer [2] and of insulin-secreting beta cells [3] took such a ranking as their starting point. A published protocol now generalises that use to target discovery [4]. In two further studies the ranking was one input among several rather than the starting point. In dentinogenesis it followed a regulator that fate mapping and regulon analysis had nominated [5]. For haematopoietic stem-cell ageing the ranking was combined with a retroviral transcription-factor screen to nominate Pbx1, and the authors then called that nominee a network hub [6]. A scan of the single-cell literature collected these and further applications of the same tools (Additional file 1: Fig. S1). In each case the list is read as a priority order, and genes near the top are the ones worth taking forward.

Two families of method are in use, and they differ in what they take as input. The zero-shot family needs only an expression matrix: a gene regulatory network is estimated from the observed cells, one gene is removed from that network, and the resulting change is propagated and ranked. scTenifoldKnk belongs to this family [7], as do CellOracle’s in silico perturbation [8] and Gene-former’s in silico deletion [9]. That tool extends the network-construction step of scTenifoldNet from differential expression to single-gene deletion [10]. A recent review surveys the family and its spatial and cross-modal extensions [11]. The supervised family is trained on measured perturbation responses and predicts the profile a perturbation would produce; scGen [12], GEARS [13] and CPA [14] represent it. Both families answer different questions on different inputs. A supervised model can return a response profile only for perturbations represented in its training data, whereas a zero-shot model is applied to atlases that contain none. Where measured perturbation data exist, the supervised route is the direct one; where they do not, the zero-shot route is what remains. Across both families we consider six perturbation methods: scTenifoldKnk, CellOracle, Geneformer, scGen, GEARS and CPA. This study evaluates the zero-shot family, on observational data, and asks what its ranking reports; supervised perturbation models are outside this comparison.

The use described above rests on an assumption stated in the tool reports but not carried into interpretation: that a gene’s rank reflects the causal consequence of removing that gene. What supports the assumption is case-level. Individual knock-outs have been reported to recover regulators that were then validated experimentally, for a regulator of cell identity [1]. Other knock-outs were supported by dependency data [2]. The same pattern appears in the tool reports and in the application papers, where a small set of named genes carries the validation. Such agreement is consistent with a causal reading without testing one, because a gene ranked high for structural reasons would produce the same agreement. Three systematic checks are absent. One asks whether the output changes when an unrelated gene is removed. A second asks whether genes with a known causal role rank above the rest. A third asks which quantity determines the output. Zero-shot evaluation of single-cell foundation models has been reported at the embedding level [15]; the knock-out ranking layer has not been evaluated in the same way. Benchmarks of network inference [16, 17] and of perturbation-response prediction [18] cover adjacent tasks, and a ranker adjusted for network topology has been proposed [19]. Whether the ranking can be corrected, or whether the causal information is absent from the input, has not been separated. Three evaluations of this family appeared the same year: eight methods across four datasets [20], a confound audit of perturbation with foundation models [21], and a benchmark under a pre-specified plan [22].

We measured this on three zero-shot methods in four configurations: scTenifoldKnk in its R and Python implementations, CellOracle 0.20.0 and Geneformer-V1-10M. The measurements cover 438 real knock-outs across nine public and three third-party datasets, 3,585 synthetic knock-outs in 36 constructed datasets, and 330 DepMap knock-outs [23, 24]. DepMap supplies an external readout of essentiality, so that a ranking can be compared with a truth it did not help to build. All quantities were computed under a criterion frozen before any result was produced. Three design elements carry the comparison. A positive control sets the causal effect and the network structure independently; two implementations of one method run under their own defaults; and the four configurations are contrasted across tool families. Six measurements follow: identity, a positive control, recovery, attribution, replication, and an alternative that costs nothing. That alternative is a detection-rate ranking, which needs no network and can be computed before one is built. The first Results subsection states the design the six share.

## 2 Results

This section states what was measured, on which data, and against which criteria. Nothing here is a result. Three zero-shot methods were run in four configurations: scTenifoldKnk in its R implementation and in its Python implementation (scTenifoldpy 0.5.1), CellOracle 0.20.0 and Geneformer-V1-10M. They were run on 438 real knock-outs in 17 runs, on 3,585 synthetic knock-outs in 36 constructed datasets, and on 330 DepMap knock-outs. The nine public datasets carry the phenomenon. Three third-party datasets test whether it survives a change of data, platform and species, at two HVG settings. Thirty-six constructed datasets form a positive control in which the causal effect is set by construction.

Three quantities were computed from each dataset’s network: output predictability (OP *R*^2^), the cross-validated *R*^2^ of a model predicting displacement from a gene’s own properties, top-*k* overlap between knock-out lists, and the zero out-degree rate. OP *R*^2^ is not by itself a test of causal content; it is used here as the quantity the criterion reports, and out-degree enters the model from the same network over which the perturbation propagates.

Table 1 lists the datasets and their knock-out counts. The first measurement needs no ground truth: it asks whether the reported list depends on which gene was removed (Fig. 1).

**Figure 1:**
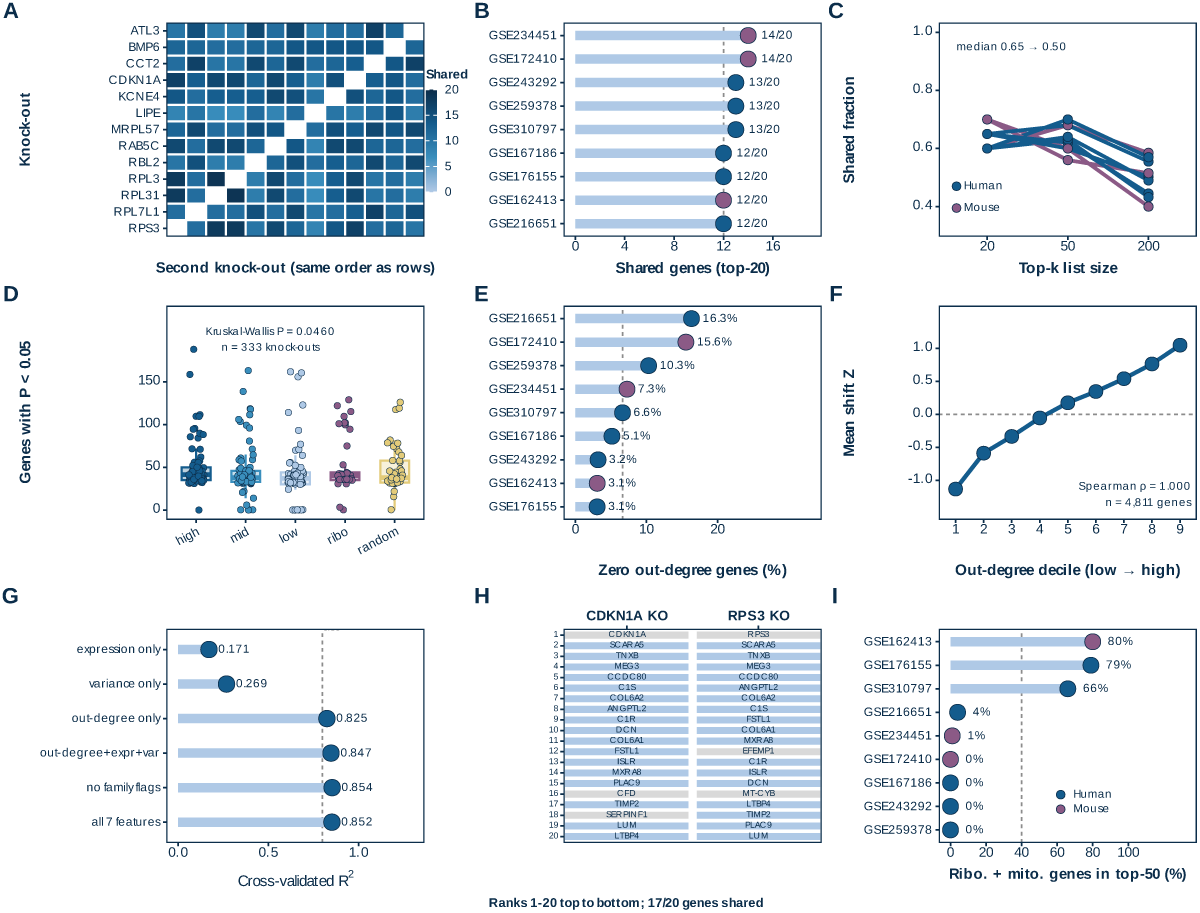
Output is independent of which gene is knocked out. (A) Top-20 overlap between every pair of the 13 knock-outs performed on GSE216651 (knee osteoarthritis synovium); cell colour is the number of shared genes, the diagonal is blank and pairwise overlap spans 7 to 19 of 20. (B) Median top-20 overlap for each of the nine public datasets (lollipop; *n* = 13 knock-outs for GSE216651 and 40 for the others), coloured by species. (C) Shared fraction (overlap/*k*) at list sizes *k* = 20, 50 and 200 for the same datasets; fractions rather than counts because *k* differs. (D) Genes with *P <* 0.05 after each knock-out, by knock-out stratum (high, mid and low mean expression, with ribosomal and random controls; *n* = 333 knock-outs pooled over datasets); boxes are median and interquartile range, points individual knock-outs. The unit of inference is the dataset; knock-outs are pooled descriptively. (E) Fraction of network genes with zero out-degree in each dataset (lollipop); the dashed line is the median across datasets. (F) Mean shift *Z* against the out-degree decile of the knocked gene (nine deciles, GSE216651; the lowest out-degree bin collapses onto the zero floor and is not drawn as a separate decile); error bars are s.e.m. and the dashed line is zero. (G) Cross-validated *R*^2^ of models predicting the displacement from features of the knocked gene, one bar per feature set. (H) Top-20 lists for two functionally unrelated knock-outs, CDKN1A and RPS3, with ranks printed in the cells. (I) Ribosomal and mitochondrial share of each dataset’s top-50 list (lollipop). Dashed lines are display boundaries fixed before the analysis (12 of 20 for list agreement; 0.80 for the OP *R*^2^ grade, OP *R*^2^ being the out-of-sample prediction *R*^2^) and were not recalibrated in this study. Statistics for every panel are given in Additional file 2.

**Table 1:**
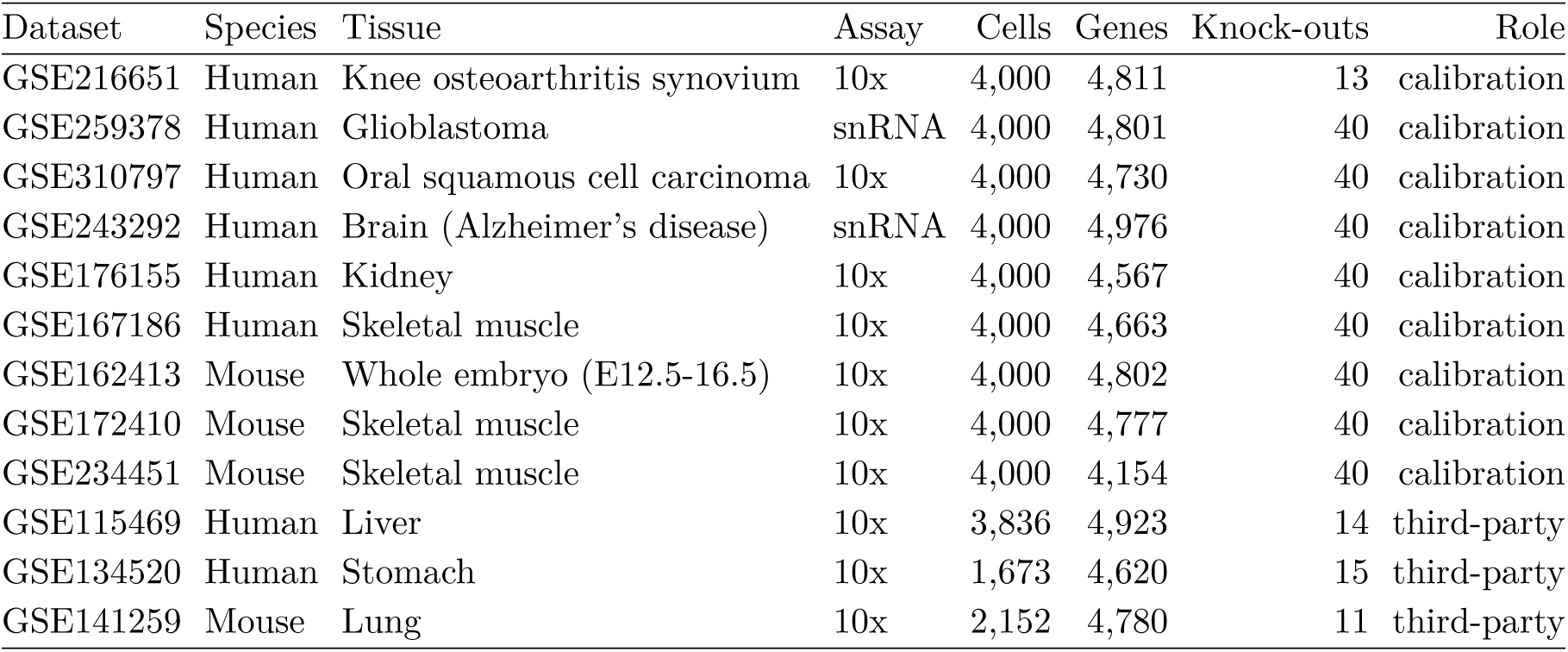
Datasets, knock-out counts and role in the argument.

| Dataset | Species | Tissue | Assay | Cells | Genes | Knock-outs | Role |
|---|---|---|---|---|---|---|---|
| GSE216651 | Human | Knee osteoarthritis synovium | 10x | 4,000 | 4,811 | 13 | calibration |
| GSE259378 | Human | Glioblastoma | snRNA | 4,000 | 4,801 | 40 | calibration |
| GSE310797 | Human | Oral squamous cell carcinoma | 10x | 4,000 | 4,730 | 40 | calibration |
| GSE243292 | Human | Brain (Alzheimer’s disease) | snRNA | 4,000 | 4,976 | 40 | calibration |
| GSE176155 | Human | Kidney | 10x | 4,000 | 4,567 | 40 | calibration |
| GSE167186 | Human | Skeletal muscle | 10x | 4,000 | 4,663 | 40 | calibration |
| GSE162413 | Mouse | Whole embryo (E12.5-16.5) | 10x | 4,000 | 4,802 | 40 | calibration |
| GSE172410 | Mouse | Skeletal muscle | 10x | 4,000 | 4,777 | 40 | calibration |
| GSE234451 | Mouse | Skeletal muscle | 10x | 4,000 | 4,154 | 40 | calibration |
| GSE115469 | Human | Liver | 10x | 3,836 | 4,923 | 14 | third-party |
| GSE134520 | Human | Stomach | 10x | 1,673 | 4,620 | 15 | third-party |
| GSE141259 | Mouse | Lung | 10x | 2,152 | 4,780 | 11 | third-party |

### 2.1 The reported genes are largely independent of which gene is knocked out

The top genes reported after a knock-out are largely the same whichever gene is knocked out. Top-20 lists of the 13 knock-outs performed on one dataset (GSE216651) were compared, then the comparison was repeated across all nine public datasets and at list sizes up to *k* = 200. Within a dataset every knock-out is scored on the same network, so overlap can be counted on a fixed gene set. Candidate predictors of displacement were the knocked gene’s out-degree, its expression, its variance and three family flags. Two independent 20-gene lists drawn from that 4,811-gene network share 0.08 genes on average, which supplies the null.

Every pair of the 13 knock-outs shared between 7 and 19 of its top 20 genes (Fig. 1a). The lowest pair, LIPE against RPL3, shared 7, and no pair shared all 20. Across the nine datasets the median overlap was 12 to 14 of 20, with a median of 13 against the expected 0.08, a 156-fold excess (Fig. 1b). The null assumes two random gene lists; a null matched on out-degree would raise the expected overlap, so the 156-fold ratio is an upper bound. The lists remained half-identical as they lengthened: the shared fraction was 0.65 at *k* = 20 and 0.50 at *k* = 200 (Fig. 1c). Duplication is therefore not confined to the top of the ranking. Overlap at this level describes near-duplicate lists, not a family resemblance among related knock-outs. The 12-of-20 line in Fig. 1b is a display criterion fixed before the analysis and not recalibrated in this study (see Methods, Evaluation criteria).

Two knock-outs with no functional relationship, CDKN1A and RPS3, share 17 of 20 genes (Fig. 1h). The size of a displacement instead tracks the gene’s own position in the network. Mean displacement rose monotonically across the nine out-degree deciles that the scale resolves, the lowest bin collapsing at the floor, and the nine decile means correlated at *ρ* = 1.000 (4,811 genes in total; Fig. 1f). Out-degree alone accounted for 0.825 of the variance in displacement, six further features raised this to 0.852, and variance or expression alone accounted for 0.269 and 0.171 (Fig. 1g). Both fits are cross-validated. The four-feature OP *R*^2^ across the nine datasets ranged from 0.8536 to 0.9902.

A gene with no outgoing edge carries no displacement by construction, because a deletion has no edge to travel along (Additional file 1: Fig. S4f). The zero out-degree rate ranged from 3.1% to 16.3% across the nine datasets (Fig. 1e), and 786 of the 4,811 genes in GSE216651 had no outgoing edge. Those genes are not a random subset: 41.0% of translation-family genes have zero out-degree. Across datasets, OP *R*^2^ falls as the zero out-degree rate rises (*ρ* = *−*0.678, *P* = 0.0448; Additional file 1: Fig. S4). Genes called significant were pooled across the 333 knock-outs, so the test describes those strata rather than inferring across datasets. The five strata differed by at most five genes (42 down to 37), and the difference was not monotone in expression; the Kruskal-Wallis *P* of 0.046 reflects this small difference (Fig. 1d). Ribosomal and mitochondrial genes reached 80%, 79% and 66% of the top 50 in the three datasets with the highest share (Fig. 1i).

The list therefore carries almost no knock-out identity while remaining highly structured. A construction in which the causal effect is known by design tests whether that structure belongs to the data or to the method.

### 2.2 A positive control with known ground truth: decoupling effect from topology changes nothing

When the causal effect is decoupled from network topology by construction, the criterion returns the same value. None of the nine public datasets contains a knock-out whose causal effect is known, so the overlap alone cannot separate a property of the data from a property of the method. The positive control asks whether the criterion can tell a construction with a causal signal from one without. Thirty-six datasets were therefore built in which the causal effect is generated by one variable and network degree by another: A1 topology-driven, A2 causal-driven and A3 mixed, 12 datasets each. The design holds the method fixed and varies only what the data contain, and both variables were verified to be decoupled. The construction and its checks are described in Additional file 1: Fig. S2.

Decoupling held in the constructions: construction degree correlated with realised effect at *ρ* = 0.043 in A2 and 0.055 in A3 (Fig. 2i). OP *R*^2^ medians were 0.916 in A1, 0.917 in A2 and 0.913 in A3, a spread of 0.004, with a Kruskal-Wallis *P* of 0.8535 (Fig. 2a, b). The three constructions are therefore indistinguishable on the quantity the criterion reports. OP *R*^2^ measures structural predictability, not causal content, and is used here only to ask whether the criterion separates the two constructions. A classifier built on OP *R*^2^ told A2 from A1 at an AUC of 0.4514 (95% CI 0.229–0.694), which did not exceed chance (Fig. 2j). The median paired difference was *−*0.00075 (95% CI *−*0.047 to 0.048), with 45.2% of resamples positive (Fig. 2k). All 36 constructions were graded RED, above the 0.80 boundary fixed before the analysis (Fig. 2l), and the three medians differ by less than the range within any single construction.

**Figure 2:**
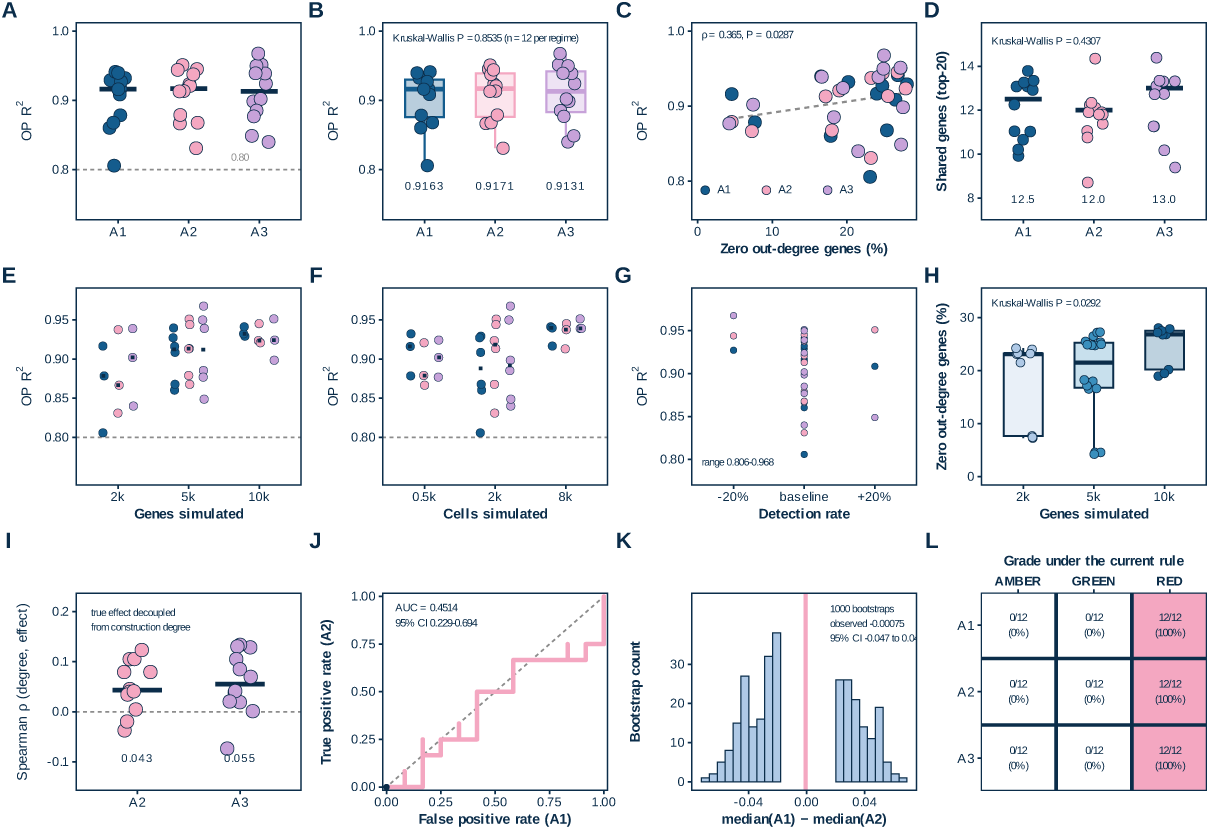
Positive control: decoupling effect from topology changes nothing. (A) OP *R*^2^ of the 36 simulated datasets by construction (A1, A2, A3; *n* = 12 each); bars are group medians. (B) The same values as box plots with points. (C) OP *R*^2^ against zero-out-degree rate in the same simulations; dashed line, least-squares fit. (D) Top-20 overlap between knock-outs within each simulation, by construction; bars are medians. (E) OP *R*^2^ by genes simulated (2k, 5k, 10k) and construction; points jittered, bars group medians. (F) OP *R*^2^ by cells simulated (0.5k, 2k, 8k), laid out as in E. (G) Paired OP *R*^2^ across the detection-rate perturbation (*−*20%, baseline, +20%); one line per simulation. (H) Zero-out-degree rate by genes simulated; boxes, median and interquartile range; points, simulations. (I) Manipulation check: Spearman correlation between the construction degree and the realised effect size, A2 and A3 only, A1 having no causal genes. (J) Empirical ROC separating A2 from A1 by OP *R*^2^ (A2 the positive class, lower OP *R*^2^ more positive); diagonal, chance; band, bootstrap interval (4,000 resamples). (K) Bootstrap distribution of the median difference between A1 and A2 (1,000 resamples); vertical line, observed difference and interval. (L) Verdict of the grading rule per simulation, rows constructions, columns grades; the empty AMBER and GREEN columns are drawn deliberately, so their absence is the result, not missing data. The 0.80 line in A, E and F is the grade boundary fixed before the analysis, not recalibrated in this study; statistics for every panel are given in Additional file 2.

The criterion is not saturated, so equality at 0.916 is not a ceiling effect. OP *R*^2^ spans 0.806 to 0.968 across the 36 constructions and a *±*20% detection-rate perturbation (Fig. 2g), and within simulations it rises with the zero out-degree rate (*ρ* = 0.365, *P* = 0.0287; Fig. 2c). The perturbation was paired, so each construction was compared with its own unperturbed version. Gene number, cell number and that perturbation left the verdict unchanged across the 3 *×* 3 grid (Fig. 2e–g; Additional file 1: Fig. S3), and the zero out-degree rate moved with gene number without moving the verdict (Fig. 2h). Top-20 lists also stayed similar within a construction: median overlap was 12.5 in A1, 12.0 in A2 and 13.0 in A3 (Kruskal-Wallis *P* = 0.4307; Fig. 2d). The overlap did not separate the three constructions either.

A construction that removes the causal signal returns the same criterion value and the same grade as one that contains it. Whether the planted causal genes can be recovered at all is a separate question. Because the criterion does not separate the two constructions, no re-ranking that uses only the wild-type network recovered the causal genes in these constructions. A re-ordering rule applied to that output would have no causal signal to reweight.

### 2.3 On data where the causal genes are known, the ranking did not recover them in the two causal constructions

On data where the causal genes are known by construction, a stronger causal effect does not raise a gene’s rank. Every gene in A2 and A3 was scored. Two questions were asked: whether the planted causal genes reach the top of the list more often than background genes, and whether a larger planted effect moves a gene further up. A2 generates the causal effect from one variable and A3 mixes that variable with network degree; the two constructions each contain 12 simulations, 24 in total. This planted signal is small relative to the network: 30 causal genes in a network of roughly 1,000 to 8,500 genes, that is 0.3% to 3%. Their summed effects run from 0.032 to 0.440, a 13.9-fold span (Fig. 3g).

**Figure 3:**
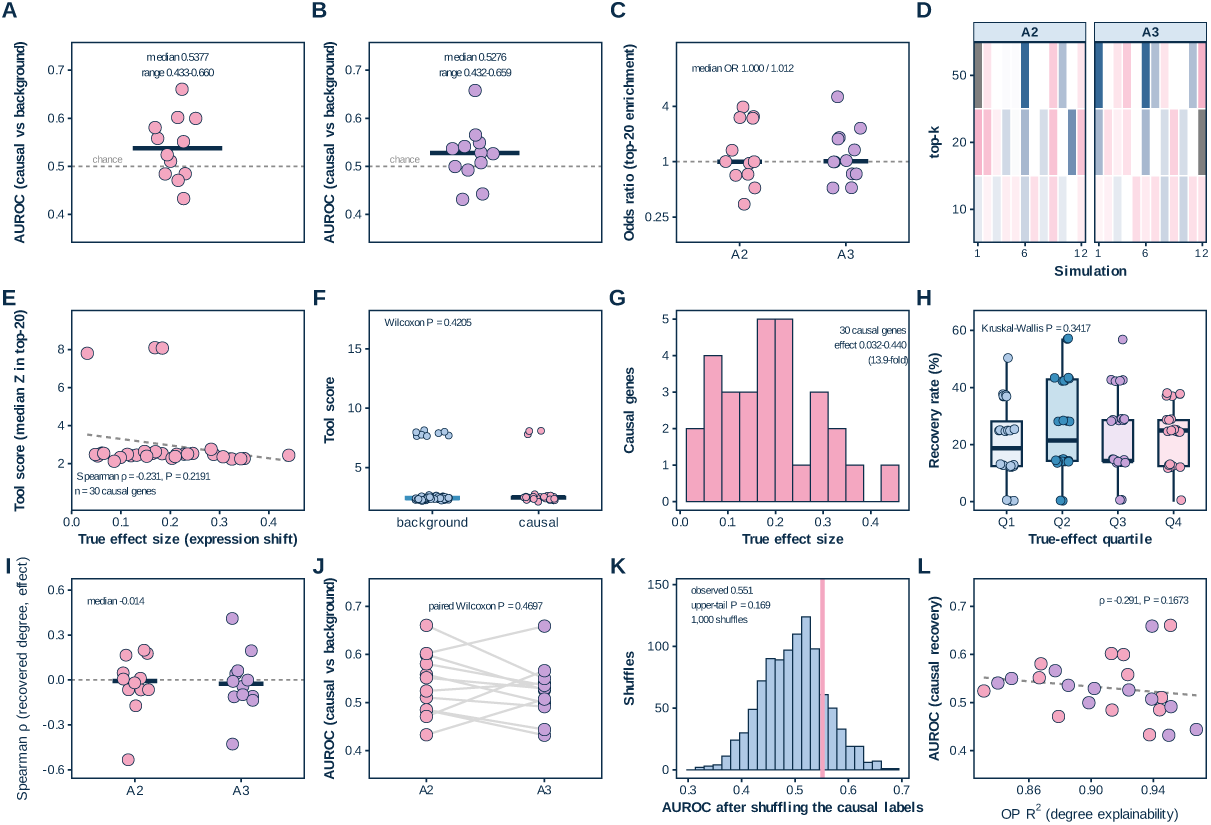
On data where the causal genes are known, the ranking did not recover them in the two causal constructions. (A) AUROC separating causal from background genes, each of the 12 A2 simulations; bar, median; dashed line, 0.5. (B) The same for the 12 A3 simulations. (C) Top-20 enrichment odds ratio (Fisher’s exact test) for the 24 A2/A3 simulations, log_2_ axis with back-transformed labels; dashed line, odds ratio of 1. (D) Top-*k* enrichment (ln odds ratio) for *k* = 10, 20 and 50, one column per simulation, faceted by construction; white, no enrichment; blue, depletion; pink, enrichment. (E) Representative A2 simulation: tool score (median *Z* of the top-20 genes) against true effect size of the 30 causal genes; dashed line, least-squares fit. (F) The same simulation: tool score of causal versus background genes; boxes, median and interquartile range; points, genes. (G) True effect sizes of the 30 causal genes, spanning 0.032 to 0.440, a 13.9-fold range. (H) Recovery rate (causal genes in the top 20) by true-effect quartile, pooled over the 24 simulations. (I) Spearman correlation between recovered out-degree and true effect within each simulation. (J) Paired AUROC in A2 versus A3 on the same simulation grid, 12 matched pairs. (K) Label-permutation null for the representative simulation (1,000 shuffles); vertical line, observed AUROC. (L) OP *R*^2^ against AUROC across the 24 simulations; dashed line, least-squares fit. The 0.5 line in A and B marks no separation and the odds-ratio line in C no enrichment; both are display references fixed before the analysis, not recalibrated here. Statistics for every panel are given in Additional file 2.

The ranking does not place the causal genes above background. Median AUROC was 0.5377 in A2 (range 0.433–0.660) and 0.5276 in A3 (0.432–0.659); each range comes from the 12 simulations of that construction, not from a confidence interval (Fig. 3a, b). AUROC was computed within each construction, with the planted genes as the positive class, and both medians sit near 0.5, the value a ranking that ignores the labels returns. At this planted effect size the ranking shows no evidence of recovery above chance. Top-20 enrichment was an odds ratio of 1.000 in A2 and 1.012 in A3 (Fig. 3c), and enrichment at *k* = 10, 20 and 50 had no consistent direction (Fig. 3d). A label-permutation null, built by shuffling effect labels within each construction, placed the observed AUROC of 0.5515 in the upper tail at *P* = 0.169 (Fig. 3k). The paired difference between A2 and A3 gave *P* = 0.4697 (Fig. 3j).

A stronger causal effect does not move a gene up. Recovery, the share of planted genes reaching the top of the list, was 0.19, 0.21, 0.14 and 0.25 across quartiles of the true effect, over the same 13.9-fold span (Kruskal-Wallis *P* = 0.342; Fig. 3h). The four quartiles cover the whole planted range, and none separates from another; the two higher quartiles do not recover more than the two lower ones. A ranking that carried even weak causal information would be expected to improve as the effect grows. The tool score was unrelated to true effect (*ρ* = *−*0.231, *P* = 0.219; Fig. 3e), and causal and background genes did not separate (Wilcoxon *P* = 0.4205; Fig. 3f). Recovered out-degree also showed no relation to effect (*ρ* = *−*0.014, a median across the 24 simulations; Fig. 3i). All three tests therefore return the same answer. Across constructions, OP *R*^2^ and AUROC correlated at *ρ* = *−*0.291 (*P* = 0.167; Fig. 3l).

A gene’s chance of recovery shows no detectable relation to how well its displacement can be predicted (Fig. 3l). What each method reports is therefore identified from the quantity its output tracks.

### 2.4 Two independent structural biases: implementation and tool family

Two implementations of one method agree only weakly, and the structural quantity behind each output differs, none of them causal. We ran both implementations of scTenifoldKnk on the same data, then related each method’s output to the structural quantities available to it. These were out-degree for scTenifoldKnk, prior target counts for CellOracle, and token coverage with six feature sets for Geneformer. The two implementations ran under their respective defaults, not a matched configuration (Methods, Perturbation methods). Fig. 4p states which methods require training perturbation data.

**Figure 4:**
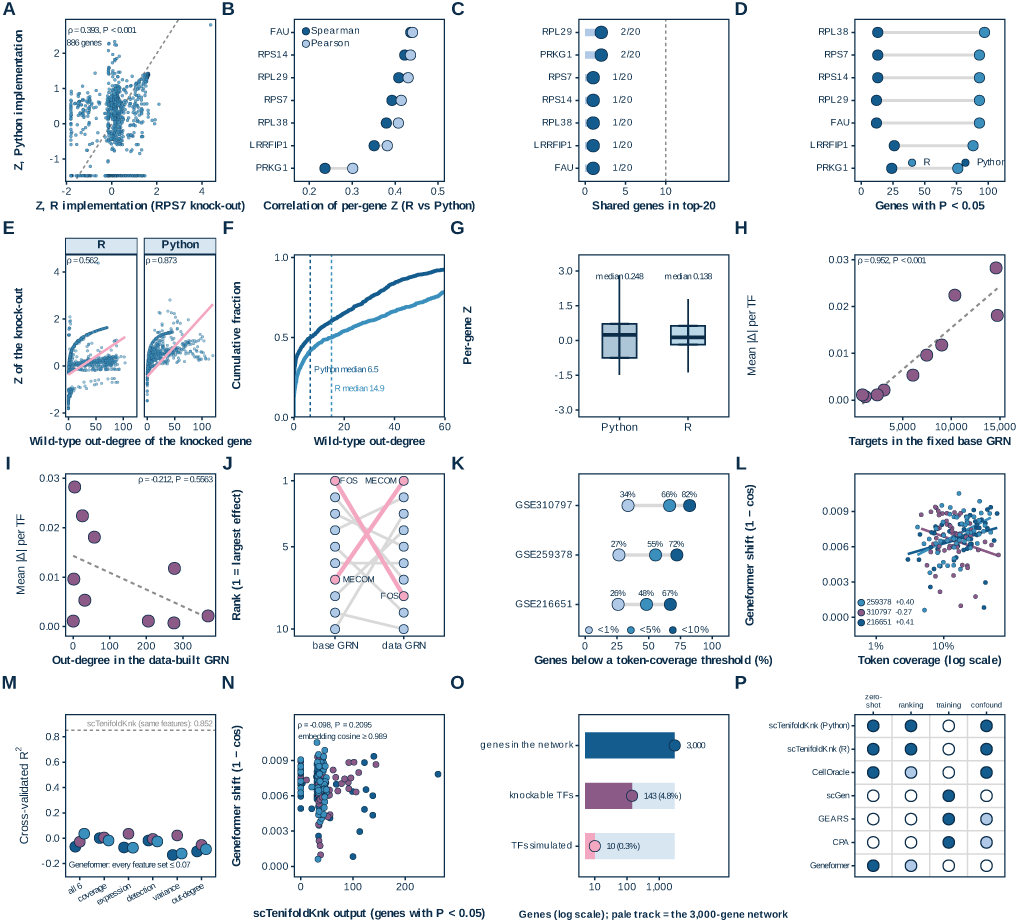
Two independent structural biases: implementation and tool family. Blocks: (A)-(D) two implementations under their own defaults; (E)-(G) output against out-degree; (H)-(J), (O) CellOracle’s prior library and knockable scope; (K)-(N) Geneformer token coverage and embedding; (P) capability of six methods across seven entries, four run here. (A) Per-gene *Z*, R against Python, for RPS7 (886 genes); dashed line, identity. (B) Per-gene rank correlation between implementations, seven knock-outs (Spearman filled, Pearson pale). (C) Top-20 overlap between implementations (lollipop); dashed line, 10 of 20. (D) Genes with *P <* 0.05 per knock-out, both implementations. (E)-(G) use that knock-out: *Z* against wild-type out-degree (E); out-degree CDF with median lines (F); per-gene *Z* distribution (G). (H) CellOracle mean |Δ| per transcription factor against target count in the fixed base GRN (10 TFs); dashed line, fit. (I) As H, against out-degree in the data-built GRN. (J) TF rank under the two definitions (slope chart). (K) Geneformer: genes below 1%, 5% and 10% token coverage, three datasets. (L) Geneformer shift (1 *−* cos) against token coverage, 167 genes, per-dataset fits; correlations differ in sign. (M) Cross-validated *R*^2^ for six feature sets predicting that shift, 55 to 57 genes; dashed line, the same model on scTenifoldKnk output (4,811 genes), a reference for another tool, not a threshold here. (N) Geneformer shift against scTenifoldKnk output, same gene. (O) Knockable scope: 143 of 3,000 genes (4.8%) are transcription factors in the base GRN and therefore knockable, and 10 were simulated. (P) Capability of six methods across seven entries against four properties (filled yes, pale partly, open no); run here: scTenifoldKnk R and Python, CellOracle, Geneformer; capability only: scGen, GEARS, CPA. Dashed criterion lines are fixed before the analysis, not recalibrated here; statistics for every panel are in Additional file 2.

Agreement between the implementations is weak gene by gene. Per-gene Z scores correlated at *ρ* = 0.393 over 886 genes (Fig. 4a), and rank correlation across seven knock-outs ranged from 0.236 to 0.436 (Fig. 4b). The two top-20 lists shared 1 to 2 genes (Fig. 4c); genes called significant numbered 76 to 97 in the R implementation and 12 to 26 in the Python implementation (Fig. 4d). That gap points the same way as the two defaults, which differ in cells per network and in quality control (see Methods, Perturbation methods). Under a matched configuration for one knock-out the correlation rose to 0.651 (971 genes), with 24 and 10 genes called significant and 3 in common, which is still far from agreement. Although both implement the same algorithm, neither reproduces the other’s gene-level output.

Within each implementation the displacement largely restates the knocked gene’s own wild-type out-degree. For the one knock-out shared by both implementations, RPS7 over 886 genes, *ρ* was 0.562 in R and 0.873 in Python (Fig. 4e). Median wild-type out-degree was 14.9 against 6.5 (Fig. 4f), and the Z medians were 0.138 and 0.248 (Fig. 4g). The strength of the out-degree relation itself depends on network density, which differs between the two runs.

CellOracle’s displacement shows no detectable relation to the knocked transcription factor’s out-degree in the data-built network (*ρ* = *−*0.212, *P* = 0.556; *n* = 10; Fig. 4i). As a consistency check, displacement scales with the number of prior targets in the fixed motif library (*ρ* = 0.952, *P <* 0.001; Fig. 4h), which is the quantity the simulator propagates along. Its ordering is not the network’s: FOS held rank 1 on displacement against 8 on out-degree, MECOM 7 against 1 (Fig. 4j). Of 3,000 genes in that network, 143 (4.8%) can be perturbed at all and 10 were simulated (0.3%; Fig. 4o).

Geneformer’s per-gene shift tracks token coverage in two datasets and reverses in the third (+0.41, *P* = 0.002; +0.40, *P* = 0.002; *−*0.27, *P* = 0.041; 55 to 57 genes per dataset; Fig. 4l). Coverage is thin: 26%, 48% and 67% of genes in GSE216651 fall below 1%, 5% and 10% (Fig. 4k). A six-feature model accounts for at most 0.074 of out-of-fold variance in 55 to 57 genes (Fig. 4m). Refitting at lower capacity leaves that ceiling in place: *−*0.067 / +0.048 / *−*0.028 at three leaves and *−*0.387 / *−*0.614 / *−*0.044 for Ridge. The shift is unrelated to scTenifoldKnk output (*ρ* = *−*0.098, *P* = 0.21; Fig. 4n), and deleting one gene moves each cell’s embedding by at most 1.1% (cosine at least 0.989). That is the scale at which this method reports. The blocks are not four versions of one bias: out-degree, prior target count and token coverage are different quantities, and each tool is bound to its own quantity, or to none that was measured.

Each tool leaves a different structural fingerprint. A change of dataset, platform and species tests whether that fingerprint belongs to these data.

### 2.5 The same pattern in third-party data, at both HVG settings, on another platform and species

Three datasets that played no part in developing the pipeline return the same structural pattern, at both HVG settings. We re-ran the identical pipeline, without retuning, on GSE115469, GSE134520 and GSE141259. Between them the three add a different sequencing platform, a second species (mouse) and a second HVG tier: each was run at 5,000 and at 10,000 highly variable genes, giving six runs. Network construction, knock-out list size and the three criteria were held at the values used for the nine public datasets. Each dataset contributes two runs, so the HVG tier is varied within a dataset and not only across datasets.

Output predictability across the six runs ranged from 0.7691 to 0.9830, with five runs graded RED and one AMBER (Fig. 5a, 5j). The zero out-degree rate ranged from 0.08% to 5.30% (Fig. 5b), and top-20 lists of different knock-outs shared 12 to 17 of 20 genes (Fig. 5c). Significant genes per knock-out numbered 31 to 76 (Fig. 5l). No run fell below the AMBER band, and reproduction here means the same three criteria returning the same qualitative verdict, not the same values.

**Figure 5:**
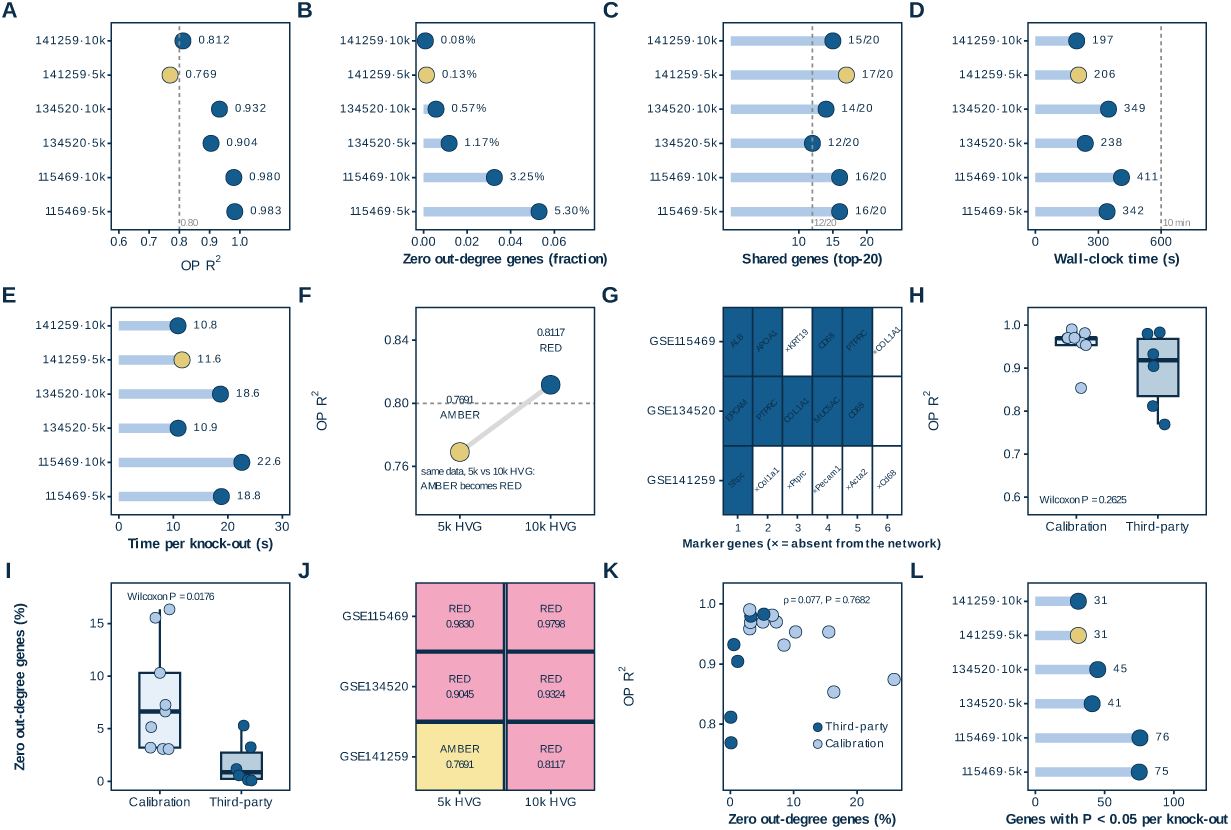
Change the dataset, the platform, the species: the same pattern, with one grade at a band boundary. (A) OP *R*^2^ of the six third-party runs (three datasets at 5k and 10k highly variable genes), lollipops; dashed line, 0.80. (B) Zero-out-degree rate of the same runs, the fraction of network genes that cannot be knocked out. (C) Top-20 overlap within each run; dashed line, 12 of 20. (D) Wall-clock time of a complete run on an otherwise idle machine; dashed line, 10-minute target. (E) Time per knock-out for the same runs, the quantity that scales with the number of genes screened. (F) GSE141259, the one run whose grade moves with the highly variable gene setting, shown at both settings on the same data; dashed line, 0.80. (G) Marker genes of each third-party dataset, name printed in the cell: deep blue, in the network and genes move; pale blue, in the network but no gene moves; white with a prefixed multiplication sign, absent from the network. (H) OP *R*^2^ of the six third-party runs against the nine public datasets; boxes, median and interquartile range; points, runs. (I) Zero-out-degree rate of the same two groups. (J) Grade of each dataset under the current rule at both highly variable gene settings. (K) OP *R*^2^ against zero-out-degree rate for all 17 tool runs, coloured by source; dashed line, least-squares fit. (L) Genes with *P <* 0.05 per knock-out in each run, median over knock-outs. The 0.80 line in A and F and the 10-minute line in D are display boundaries fixed before the analysis, not recalibrated here; 12 of 20 is the list-agreement criterion fixed before the analysis. Statistics for every panel are given in Additional file 2.

Against the nine public datasets the third-party runs are not different in output predictability (*P* = 0.2625; Fig. 5h). They differ in the zero out-degree rate (*P* = 0.0176; Fig. 5i), which is lower here. One verdict moves with the HVG tier: GSE141259 graded 0.7691 at 5,000 genes and 0.8117 at 10,000, and the lower of those two is the one AMBER case (Fig. 5f). Across all 17 runs, output predictability and the zero out-degree rate are unrelated (*ρ* = 0.077, *P* = 0.768; Fig. 5k). That correlation is computed across runs, not across datasets within a run.

Marker recovery is partial and is reported as presence in the network. Seven of 17 markers were absent, and five of six mouse markers for GSE141259 were queried as human symbols (Fig. 5g). Runs were also short, at 197 to 411 s per run and 10.8 to 22.6 s per knock-out (Fig. 5d, 5e).

That excludes an explanation in which the pattern belongs only to the nine public datasets. Whether anything that costs nothing does better is a separate question.

### 2.6 A zero-cost baseline: detection rate matches or exceeds virtual knock-out on the same genes

Detection rate, a baseline that needs no network, ordered the same genes above virtual knock-out in five of six datasets. The paired gain was positive in 97.6% of 1,000 resamples (+0.129, 95% CI 0.0017–0.2543; Fig. 6e, 6i). On the 330 DepMap knock-outs we ranked the same genes by the tool’s score, by mean expression, by detection rate, and by out-degree. Each ranking was scored against Chronos essentiality; 45 of the 330 genes are essential. Out-degree, the quantity that accounts for most of the tool’s output, reached AUROC 0.363 (95% CI 0.278–0.453; Additional file 2): the structural quantity itself is not an essentiality ranking; its interval lies below 0.5.

**Figure 6:**
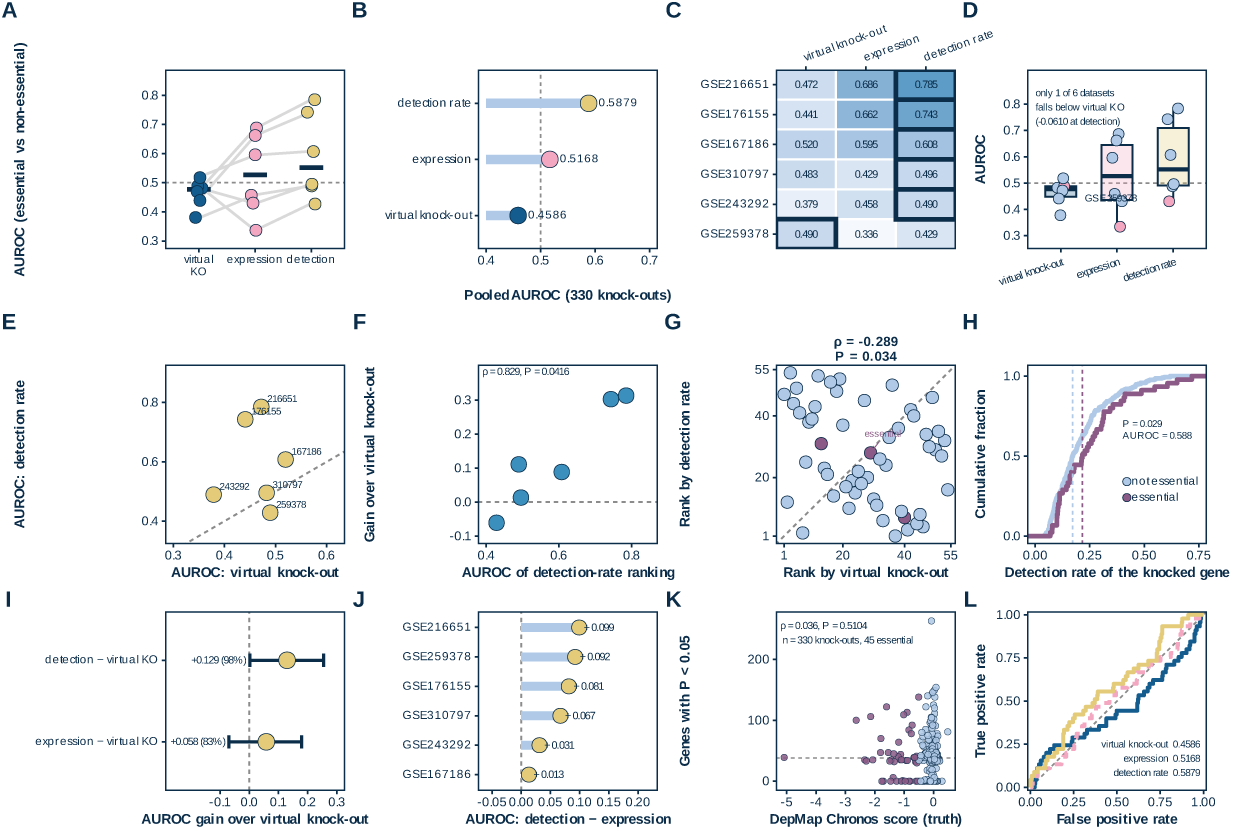
If not this tool, then what: a usable alternative. Virtual knock-out throughout is scTenifoldKnk, Python implementation (scTenifoldpy 0.5.1). (A) AUROC separating essential from non-essential genes for three rankings (virtual knock-out, expression, detection rate) in six datasets; grey lines join one dataset, points jittered, bars medians. (B) Pooled AUROC over the 330 DepMap knock-outs for the three rankings, values printed at the bar ends. (C) AUROC of the three rankings per dataset (heatmap, darker higher); black outline, best ranking of each row. (D) AUROC distribution across the six datasets for the three rankings; boxes, median and interquartile range; GSE259378 highlighted. (E) Paired AUROC of virtual knock-out and detection rate, one point per dataset; dashed diagonal, identity. (F) Gain of the detection-rate ranking over virtual knock-out against the detection-rate AUROC; dashed line, least-squares fit. (G) Rank by virtual knock-out against rank by detection rate for GSE167186 (54 knock-outs); dashed diagonal, identity; essential genes mauve. (H) Cumulative distribution of detection rate for the 330 DepMap knock-out genes, essential (*n* = 45) versus non-essential (*n* = 285); dashed lines, group medians. (I) AUROC gain over virtual knock-out with 95% bootstrap intervals for the two alternative rankings; the interval for the expression ranking includes zero. (J) Per-dataset difference in AUROC between detection rate and expression. (K) Chronos score against genes with *P <* 0.05 for all 330 knock-outs, essential genes mauve; dashed line, median significant-gene count, not a threshold. (L) Pooled ROC curves for the three rankings, AUROCs printed in the panel; dashed diagonal, chance. Dashed criterion lines are fixed before the analysis, not recalibrated here; statistics for every panel are given in Additional file 2.

Pooled AUROC over the 330 knock-outs was 0.459 for scTenifoldKnk, Python implementation (scTenifoldpy 0.5.1), 0.517 (95% CI 0.423–0.608) for expression and 0.588 (95% CI 0.497–0.677) for detection rate (Fig. 6b, 6l; Additional file 2). The tool’s estimate of 0.459 is not distinguishable from chance (95% CI 0.357–0.566). Two further readouts of the same displacement vector do not move it: the L2 norm gives 0.541 (0.447–0.636) and max*|Z|* gives 0.510 (0.413–0.611). Across these readouts the tool’s ranking remained not distinguishable from chance. Within tertiles of an out-degree-stratified panel of 530 knock-outs (the 330 analysed above plus 200 additional stratified knock-outs; 97 essential), AUROC was 0.431, 0.492 and 0.496 (Additional file 2). Each stratum contains 30 to 34 essential genes, so these values are descriptive; within this frame the pooled result was not explained by out-degree.

The exception in that comparison was GSE259378 at *−*0.061 (Fig. 6e), and the gain was computed on the genes for which the tool returns an output. Gain and baseline strength moved together across the six datasets (*ρ* = 0.829, *P* = 0.042; Fig. 6f). The expression baseline is reported once, as a second zero-cost direction, with a gain of +0.0582 (Fig. 6i).

Detection rate was higher for essential than for non-essential genes, 0.217 against 0.172 (one-sided *P* = 0.029; two-sided *≈*0.058; Fig. 6h). The number of genes called significant after knock-out showed no such difference (one-sided *P* = 0.815), and against Chronos that hit count gave *ρ* = 0.036 (*P* = 0.510; Fig. 6k). In GSE167186 the detection-rate ranking was inversely related to the virtual knock-out ranking (rank–rank *ρ* = *−*0.289, *P* = 0.034; Fig. 6g). Detection rate exceeded expression in every dataset, by 0.013 to 0.099 (Fig. 6j). Per-dataset AUROC for the three rankings is given in Fig. 6a and 6c, and the distribution across datasets in Fig. 6d.

AUROC describes the full ranking; the paired gain is not a claim about the top few genes. Sixty knock-outs took 1,856 s, against 302 s to read out-degree from the same networks, a 6.1-fold difference (Additional file 1: Fig. S5). At the level of a single knock-out the ratio is 25,900-fold, because the network is built once. The ranking the tool returns orders genes by a property they hold in the network, and on this panel that property did not track essentiality.

A ranking that costs nothing was closer to external essentiality than the tool was. What the expensive ranking still reports is a separate question, and it has an answer.

## 3 Discussion

The ranking reports the knocked gene’s position in a network, fixed before any knock-out is simulated: edges are estimated from the cells that supply the expression matrix, or from a fixed cis-regulatory prior [16, 25], so every edge predates every deletion. A deletion propagates along outgoing edges, so displacement measures the reach of the deletion through the graph, not its effect on the cell.

Out-degree alone accounts for 0.825 of the variance in displacement in the GSE216651 network, and seven features together reach 0.852 (Fig. 1g). Both implementations of scTenifoldKnk tie displacement to the same quantity, at *ρ* = 0.562 for the R run and 0.873 for the Python run, for the single knock-out the two share (Fig. 4e).

CellOracle reaches the same end through a fixed prior. Only transcription factors in that library can be perturbed, 143 of 3,000 genes (4.8%), and 10 were simulated (Fig. 4o). Its displacement shows no detectable relation to out-degree in the data-built network (*ρ* = *−*0.212, *P* = 0.556; Fig. 4i). Against the target count of the same factor in the prior library it scales with that count (*ρ* = 0.952; Fig. 4h). Geneformer shifts each cell’s embedding by at most 1.1% (cosine at least 0.989; Fig. 4n), the scale at which that method reports.

The list ranks structural importance, a property measurable from the network itself. Degree distributions in these graphs are heavy-tailed [26], and centrality in protein networks was tied to essentiality [27]; these concern undirected protein-interaction degree, not directed out-degree. That tie is weaker than it first appeared: it shrinks once interaction sampling is corrected [28], essentiality tracks module membership more closely than degree [29], and the explanation did not survive scrutiny of network properties [30]. A ranking built on out-degree therefore restates a quantity whose link to essentiality is weak and contested. On this panel that quantity ranked essential genes lower, not higher (AUROC 0.363, 95% CI 0.278–0.453; Additional file 2); it is a different object from a causal priority list.

Two quantities make the reading checkable before ground truth exists: the share of network genes with no outgoing edge, 3.1% to 16.3% across the nine datasets (Fig. 1e), and OP *R*^2^, 0.8536 to 0.9902 (Additional file 2). Sparse counts contribute, and their depth dependence is measurable [31]. Zeros in expression also carry biological signal [32], but we did not partition that share into technical and biological components. Where OP *R*^2^ is high, the ordering comes mostly from the genes’ attributes.

Three uses follow. Hub candidates can be nominated from the top of the list, where genes of high out-degree collect, once the ribosomal and mitochondrial genes that fill 66 to 80% of the top 50 are removed (Fig. 1i). Hypotheses about network position can be stated so a targeted experiment can test them. Pathway-level filtering can narrow a gene set before any perturbation is run. Each use asks what a gene is in the network; none asks what the cell does without it.

Three evaluations published the same year converge [20–22]: six of eight methods did not detect the transcription-factor-to-pathway signal, and CellOracle’s perturbation direction agreed with CRISPRi in 40.9% of cases, as often as chance [20]. None of 46 pre-specified tests met its criterion, knock-out displacement did not exceed expression-matched control genes, and a naive predictor did as well [22]. Not one-sided: on K562 under a CRISPRa protocol, Geneformer V2 showed a detectable correspondence between perturbation direction and effect size (*ρ* = 0.30) [22]. That is V2 in a signed task; V1 was asked here to order genes, so the two do not conflict. The addition here is attribution: each output traced to one structural quantity, out-degree for both scTenifoldKnk implementations, prior target count for CellOracle, none for Geneformer; a 36-construction positive control; 330 DepMap knock-outs as external truth; and two implementations of one method that disagree (*ρ* = 0.393, 1 to 2 genes of 20 shared). The three evaluations did none of the four. Earlier comparisons were narrower: linear baselines were competitive [18], generalisation is benchmarked elsewhere [33–35], and the tool’s report caveated a bias toward regulatory genes [7].

### When a virtual knock-out ranking can be read, and how

A list can be read once two things are stated: what it measures, and how much of its ordering the network accounts for. Three rules follow.

1. Report the zero out-degree rate before fitting; it needs no model and shows which part of the network a knock-out cannot reach. Compute OP *R*^2^ only if that rate is low and structure is the suspected driver. A value above 0.80 is reported as above the display boundary fixed before the analysis.
2. Read the top of the list as structural importance, and only after those families are removed: hub candidates, hypotheses about network position, pathway-level filtering. The unfiltered list is not a hub list, nor a priority order for causal testing.
3. Start from detection rate when genes must be ordered by necessity. Essentiality is not selectivity, so this ranking orders experiments rather than nominating targets. On the same 330 genes it reached 0.588 (95% CI 0.497–0.677) against 0.459 (95% CI 0.357–0.566), a paired gain of +0.129 (95% CI 0.0017–0.2543; Fig. 6b, 6l). That answers an ordering question, not a causal one.

A topology-adjusted ranker (TAKO) [19] was untested; on scTenifoldKnk the ranking did not recover planted causal genes. Thirty-six decoupled constructions graded RED, and separating causal from topology-driven ones gave an AUC of 0.4514 (Fig. 2j, 2l): the criterion read the same value with or without a causal signal. AUROC for recovery was 0.5377 in A2 and 0.5276 in A3, near the value a ranking that ignores the labels returns (Fig. 3a, 3b). On external essentiality the tool’s ranking gave 0.459 (95% CI 0.357–0.566) against 0.588 (95% CI 0.497–0.677) for detection rate (Fig. 6b). A re-ordering rule applied to that output has no causal signal to reweight. Adversarial validation of Geneformer deletions controls single-gene claims with an expression-matched null [36], a different unit from the ranking. Where a causal reading is required, the ranking must be replaced by interventional or genetic evidence: pooled perturbation screens, or germline variation under a causal design. Detection rate only orders the sequence of tests.

### 3.1 Limitations

1. The simulator does not reproduce every statistic of the real data (Additional file 1: Fig. S2), so the synthetic arm tests only whether the criterion can distinguish constructions.
2. The causal-recovery analysis uses 12 versus 12 simulated datasets per comparison, so AUROC is reported as a range across simulations rather than as a confidence interval.
3. All analyses are computational; no wet-lab validation is presented.
4. Four configurations of three perturbation methods were executed; the supervised family (scGen, GEARS, CPA) was characterised on capability but not run on these data.
5. The expression baseline has a 95% confidence interval that crosses zero.
6. Geneformer’s cross-validated *R*^2^ of at most 0.074 is estimated in 55 to 57 genes per dataset; it means that the six measured structural features do not account for the output, not that no driver exists.
7. CellOracle’s correlation with data-derived out-degree rests on ten transcription factors.
8. The external arm contains three datasets and six runs.
9. The analysis addresses single-gene knock-out followed by downstream ranking; it does not test compound, chemical or perturbational-profile prediction.
10. Corrections for multiple comparisons were not applied; nominal *P* values are reported throughout.

## 4 Conclusions

For the two network-based methods tested, scTenifoldKnk and CellOracle, the ranking reports a structural property of the gene, not the consequence of removing it. That property differs by method: out-degree for scTenifoldKnk, in both implementations, and prior target counts for CellOracle. For Geneformer, no measured structural feature accounted for more than a small fraction of the shift. The same pattern returned in three datasets that played no part in the pipeline, at both highly variable gene settings, on another platform and in another species. Against external essentiality, the scTenifoldKnk Python implementation (scTenifoldpy 0.5.1) reached AUROC 0.459 (95% CI 0.357–0.566), not distinguishable from chance. On data where the causal genes were planted, that ranking showed no evidence of recovery above chance at that planted effect size, in either construction (median AUROC 0.538 and 0.528). Decoupling the causal effect from topology left the criterion value and the grade unchanged, and re-ranking the same wild-type network did not recover the causal genes in the two causal constructions. A detection-rate ranking, which costs no network, reached 0.588 (95% CI 0.497–0.677) on the same genes; its gain over virtual knock-out was +0.129 (95% CI 0.0017–0.2543; 97.6% of resamples positive). Every list should be read with its zero out-degree rate and OP *R*^2^ reported alongside it. Where a causal reading is required, interventional or genetic evidence should be used, and a detection-rate ranking only orders the sequence of tests.

## 5 Methods

### 5.1 Datasets and knock-out design

Nine public single-cell datasets were analysed, six human and three mouse, generated on the 10x Chromium platform or by single-nucleus RNA sequencing (Table 1). The human data span knee synovium, glioblastoma, oral squamous cell carcinoma, Alzheimer’s disease brain, kidney and skeletal muscle. Mouse data span whole embryo and skeletal muscle; no two datasets share cells.

Each dataset was exported at 4,000 cells and reduced to 5,000 highly variable genes, taken as the genes of highest raw-count variance (random seed 1); no mean–variance-stabilised selection was applied. Within the pipeline, cells with a library size below 1,000 counts or a mitochondrial fraction above 0.1 were removed, genes detected in fewer than 5% of cells (min_percent = 0.05) were dropped, and 4,154 to 4,976 genes remained in the reconstructed network. A PCNet was built from 2,000 cells drawn at random from the exported matrix, and every dataset was rebuilt 10 to 20 times. The exported matrix was fixed per dataset, so every method saw the same cells and the same genes.

GSE216651 was knocked out at 13 genes; the other eight datasets at 40 genes each, for 333 knockouts across the nine datasets. Knock-out genes were drawn from tertiles of mean expression, one gene per tertile for GSE216651 and ten for the others, with five ribosomal and five random controls added in every dataset.

Three datasets outside this set were processed twice, at 5,000 and at 10,000 highly variable genes: GSE115469 (liver), GSE134520 (stomach) and GSE141259 (lung). The six runs knocked out 11 to 15 genes each, 79 in total, chosen by the same rule as the main datasets. Two further GSE216651 runs, at 2,000 and at 10,000 highly variable genes, added 13 knock-outs each, so the 17 runs contributed 438 real knock-outs.

Thirty-six synthetic datasets were generated, twelve for each of three constructions. In A1 the effect of a knock-out is driven by network position; in A2 by a planted causal effect; A3 mixes the two. They contributed 3,585 knock-outs (1,196, 1,193 and 1,196). Each construction covered a 3 *×* 3 grid of 2,000, 5,000 and 10,000 genes by 500, 2,000 and 8,000 cells, to which a seed-2 replicate and two *±*20% detection-rate variants were added, so the centre cell (5,000 genes, 2,000 cells) carries four of the twelve and the other eight cells one each.

Counts were generated by a Gaussian copula with group-specific zero-inflated negative binomial margins, calibrated on the GSE216651 wild-type network and expression pool: a configuration-model graph carrying the observed out-degree sequence and zero out-degree fraction, edge weights drawn from the observed distribution, and a precision matrix rescaled to spectral radius 0.80. Marker structure came from 18 cell groups with logit-normal gene loadings (*µ* = 1.0, *σ* = 1.2), 16% of genes set near zero, and per-group modulation of detection and mean (*κ* 1.6 and 1.0). In A2 and A3 the planted causal genes were given module sizes and effect sizes, and the assignment was permuted 800 times until construction degree and effect size were decoupled (*|ρ| <* 0.10). Each simulation knocked out 100 genes: eight per construction-degree decile plus random draws in A1, and the 30 causal and 15 hub genes plus random draws in A2 and A3.

External essentiality was taken from DepMap CRISPRGeneEffect 26Q1, scored with Chronos, in which more negative values indicate stronger essentiality: 330 knock-outs over the six human datasets, 45 of them essential at Chronos *< −*0.5. The release covers 18,531 genes across 1,208 cell lines. Those 330 knock-outs were drawn uniformly at random from the genes passing quality control, one panel per human dataset, and intersected with DepMap. This retains the natural expression–essentiality association (observed *r* = 0.017 in the sample), which the expression-stratified panel removes. The three AUROC values therefore describe this panel. An out-degree-stratified panel of 530 knock-outs (the 330 above plus 200 additional stratified knockouts; 97 essential; Additional file 2) was used to check that the pooled result was not explained by out-degree.

Dataset composition, knock-out counts and network counts are listed in Table 1; both highly variable gene settings are in Additional file 1: Fig. S6.

### 5.2 Perturbation methods

Three methods were executed in four configurations: scTenifoldKnk in its R and Python implementations, CellOracle 0.20.0 and Geneformer-V1-10M.

Both scTenifoldKnk implementations rebuild an ensemble of PCNets, decompose the ensemble as a tensor, align the knock-out manifold onto the wild-type manifold, and return a per-gene displacement *Z* for every network gene [7, 10]. A PCNet is a principal-component regression network fitted to a random subset of cells, so the ensemble reduces the influence of which cells were drawn. The per-gene difference in the aligned space is the displacement, and its package *P* value compares the gene against the rest of the network.

Ensembles were built 10 to 20 times per dataset, with 2,000 cells per PCNet, three components, *q* = 0.95, 30 manifold dimensions and seed 1. The network was read once per dataset and reused for every knock-out in that dataset. Reading the network took 302 s; sixty knock-outs on the same network took 1,856 s.

The two implementations differed in their defaults for cell handling: R ran with QC disabled and up to 500 cells per PCNet (10 PCNets), while Python applied its default QC, retaining 260 of 1,000 cells and sampling from those. All other parameters (10 networks, 3 components, *q* = 0.95, 30 manifold dimensions, seed 1) were matched.

CellOracle 0.20.0 combined a fixed base GRN — the gimmemotifsv5 parquet at false-positive rate 2% and motif threshold 10, 1,094 transcription-factor columns — with a per-cluster GRN inferred from the same data [8]. Clustering used *n_neighbors_*= 15 and Leiden resolution 0.25; the GRN used knn_imputation(n_pca_dims = 30, balanced = True) and get_links(alpha = 10, bagging_number = 5), with n_jobs = 8. CellOracle ran on GSE216651 only. Its in silico perturbation could target only transcription factors present in the base GRN: 143 of 3,000 network genes (4.8%), ten of which were simulated, because a perturbation of a gene outside the base GRN returns a zero shift by construction. The output was the simulated expression shift and its projection into the embedding.

Geneformer-V1-10M, the 10-million-parameter checkpoint, performed in silico deletion per dataset with model_version = V1, max_ncells = 2,000, forward_batch_size = 32, emb_layer = 0, emb_mode = cell and nproc = 1 [9]. Each deleted gene was evaluated 55 to 57 times, and the output was one minus the cosine similarity between the deleted-cell and wild-type embeddings. Its displacement was compared against scTenifoldKnk output for the same genes.

The same knock-out gene list was used for every method within a dataset, so rankings could be compared gene by gene. scGen and GEARS were not run on these data [12, 13]. CPA was assessed from its published description [14]. The three appear only in the capability matrix of Fig. 4p.

### 5.3 Evaluation criteria

Output predictability (OP *R*^2^) is the cross-validated coefficient of determination of a model that predicts a gene’s displacement from properties of that gene [7]. Two feature sets are in use here and are not interchangeable. The seven-feature set adds three family flags (ribosomal, mitochondrial, translation) to log out-degree, log mean expression, detection rate and log variance, and is the fit behind Fig. 1g. The deposited tool, the per-dataset panel and the 36 simulations use the four features without the flags, and the panel computed this way spans 0.8536 to 0.9902 across the nine datasets. Because the arms differ in feature count, their values are compared only on whether each exceeds 0.80, not on magnitude.

Fits used five-fold cross-validation and a histogram gradient-boosting regressor with 200 iterations, a learning rate of 0.05 and at most 31 leaves per tree. The target was the gene’s mean displacement *Z* across that dataset’s knock-outs, not a single knock-out’s vector, and OP *R*^2^ was computed per dataset from those outputs. Single-feature refits on out-degree, expression and variance used the same folds.

Top-*k* overlap is the size of the intersection of two ranked lists; the shared fraction is that size divided by *k*. Both were computed at *k* = 20 and *k* = 200. The zero out-degree rate is the fraction of network genes with no outgoing edge in the averaged PCNet. It was computed for every run before any other criterion, because such a gene cannot propagate a perturbation.

Three grades were fixed before any result was produced: RED above 0.80, AMBER from 0.50 to 0.80, and GREEN below 0.50. No threshold was revised during the study. A run was graded on its OP *R*^2^ alone, without reference to its AUROC, so that grading and causal accuracy could be compared as separate quantities.

The 12-of-20 overlap screen comes from the same pre-specified criteria and is a display criterion, not a null-based cut. Two random 20-gene lists drawn from a 4,811-gene network overlap by 0.08 genes on expectation, against which the screen is read.

The knockable scope of a method is the set of genes it can perturb in principle. Two rankings that cost no network were computed on the same genes: detection rate, the fraction of cells in which a gene is detected, and mean expression, the average count of that gene. Both used the exported matrix behind the networks. Mean expression was the pre-specified comparator; detection rate was added after that comparison returned an interval spanning zero (+0.058, 95% CI *−*0.069 to 0.179). Both are reported.

For each DepMap knock-out the virtual knock-out score was the number of genes called significant at *P <* 0.05. The same displacement vector was also scored by its L2 norm and by its maximum *|Z|*. These three readouts gave AUROC 0.459 (95% CI 0.357–0.566), 0.541 (0.447–0.636) and 0.510 (0.413–0.611). Essentiality thresholds of Chronos *< −*0.3 and *< −*0.7 gave 0.437 and 0.478. The ground truth for these scores was the DepMap Chronos score [23, 24].

### 5.4 Statistical analysis

Tests were chosen by design: Kruskal-Wallis for more than two groups, Wilcoxon for two, paired where the same genes or simulated grid points were compared and unpaired otherwise.

Correlations are Spearman coefficients across genes within a dataset unless a cross-dataset relation is named. AUROC was computed by a rank method on the pooled panel, and per dataset with the same function.

Confidence intervals came from resampling: 4,000 bootstrap draws for the pooled AUROC values, and 1,000 for median differences and AUROC gains. The per-construction causal-recovery AUROC is reported as a range across simulations, because 12 datasets per construction do not support an interval. Resampling was over knock-out rows for the DepMap panel and over datasets for cross-dataset contrasts. Enrichment of essential genes within a list used Fisher’s exact test. Label permutations shuffled knock-out labels within a dataset 1,000 times and recomputed the statistic, giving an upper-tail *P*; a 30-permutation label null was computed for the real datasets. Every interval quoted in the text carries the resample count of its estimate.

The unit of analysis was the dataset or the simulation, not the cell, so that cell number could not inflate a *P* value. Paired comparisons between the A2 and A3 constructions were matched on grid point rather than on simulation identifier.

Significance was nominal *P <* 0.05 throughout, and Wilcoxon tests were two-sided unless stated.

Where the two scTenifoldKnk implementations were compared, genes were matched on identifier and restricted to the 886 genes shared by both runs. Cross-dataset statements use the median over datasets, with the range over datasets given beside it. A Kruskal-Wallis test was followed by pairwise Wilcoxon only where the omnibus test is named in the text. Every interval was recomputed from the stored per-knock-out tables rather than read from a figure.

The essential-versus-non-essential comparison was pre-specified as one-sided; the two-sided value is *≈*0.058 and is reported alongside.

### 5.5 Software and data availability

Analyses ran in Python 3.10 with scTenifoldpy 0.5.1, CellOracle 0.20.0 and Geneformer-V1-10M, using numpy, scipy and scikit-learn. The R implementation of scTenifoldKnk was run from its CRAN release under R 4.5.2.

Random seeds were 1 for network construction and manifold alignment, and were fixed per analysis for bootstrap and permutation.

The nine datasets used for the main analysis are available from GEO under GSE216651, GSE259378, GSE310797, GSE243292, GSE176155, GSE167186, GSE162413, GSE172410 and GSE234451. GSE115469, GSE134520 and GSE141259 are the three further datasets. DepMap CRISPRGeneEffect 26Q1 was obtained from the DepMap portal.

The three further datasets were taken from GEO with their original supplementary files and exported under the same pipeline; they retain 3,836, 1,673 and 2,152 cells. DepMap 26Q1 files were used as distributed, without re-scoring.

All analysis scripts and the figure scripts are provided as Additional file 3; the summary tables behind every figure are provided as Additional file 2. Resource footprints are given in Additional file 1: Fig. S6.

The summary tables in Additional file 2 carry the per-panel values, so any quoted value can be recomputed from them.

## Supporting information

Per-panel statistics and source tables

Source code for all analyses and figures

## Declarations

### Ethics approval and consent to participate

Not applicable. The study reanalysed publicly available, de-identified datasets deposited in GEO and DepMap; no new human participants, human tissue or animals were involved.

### Consent for publication

Not applicable.

### Availability of data and materials

The nine single-cell datasets analysed in this study are available in the GEO repository under accessions GSE216651, GSE259378, GSE310797, GSE243292, GSE176155, GSE167186, GSE162413, GSE172410 and GSE234451. The three further datasets are available under GSE115469, GSE134520 and GSE141259. DepMap CRISPRGeneEffect 26Q1 is available from the DepMap portal. All analysis scripts are provided as Additional file 3, and the summary tables behind every figure as Additional file 2.

### Competing interests

The authors declare that they have no competing interests.

## Funding

The authors did not receive any specific grant from funding agencies in the public, commercial, or not-for-profit sectors.

## Authors’ contributions

Jiale Cui, Meixuan Li, Rui Zhou, Haoyu Zhang, Saihang Xu, Lei Li and Rong Zhang contributed to the study design, the analysis and the interpretation of the data. Jiale Cui wrote the first draft; Lei Li and Rong Zhang supervised the project and revised the manuscript. All authors read and approved the final manuscript.

## Acknowledgements

Not applicable.

## List of abbreviations

OP R2: output predictability, the cross-validated coefficient of determination of a model predicting a gene’s displacement from that gene’s properties
AUC: area under the curve
AUROC: area under the receiver operating characteristic curve
CI: confidence interval
GRN: gene regulatory network
HVG: highly variable gene
PCNet: principal-component network
TF: transcription factor
KO: knock-out
DepMap: Cancer Dependency Map
Chronos: cell population dynamics model used to score gene fitness effects in DepMap
snRNA: single-nucleus RNA sequencing
UMAP: uniform manifold approximation and projection
GEO: Gene Expression Omnibus
QC: quality control

## Additional files

**Additional file 1** Supplementary figures S1–S6 (PDF).

**Additional file 2** Per-panel statistics and the base tables behind every figure (XLSX).

**Additional file 3** Source code for all analyses and figures (ZIP).

## Additional file 1: Supplementary figure legends

### Additional file 1: Fig. S1

**Fig. S1 | The literature pool and the genes it nominates.** (A) Records retrieved by each of the three search strings (lollipop): the generic virtual knock-out string, scTenifold and Geneformer. (B) Share of those records that carry an abstract. (C) Cumulative records by publication year for the three strings, one point per year. (D) Extraction funnel: parsed full texts, unique gene symbols and genes present in DepMap. (E) Number of papers naming each extracted gene (histogram, bin width 1). (F) The twelve most frequently nominated genes (lollipop). (G) Records per year with all three strings pooled (bars). (H) Cumulative share of gene mentions as a function of gene rank (log x axis). (I) Papers naming each gene against gene rank (log x axis), showing the long tail of single-mention genes. Counts for every panel are given in Additional file 2.

### Additional file 1: Fig. S2

**Fig. S2 | Simulator fidelity and the fit gate.** (A) Kolmogorov-Smirnov distance between simulated and real degree distributions for all 37 simulated datasets (histogram); the dashed line is the 0.10 tolerance. (B) Simulated against real median out-degree, one point per simulation, coloured by gene tier; the dashed line is identity. (C) Simulated against real zero-out-degree rate; the dashed line is identity. (D) Simulated against real mean-variance slope; the dashed line is identity. (E) Simulated against real mean-variance *R*^2^; the dashed line is identity. (F) Median detection rate, simulated versus real, by gene tier (paired bars). (G) Structural specificity, simulated versus real, by gene tier. (H) Distribution of gene and cell retention after quality control (overlapping histograms). (I) Fraction of the 37 simulations passing each fidelity criterion, shown separately and all together. The 37 are the 36 of the 3 *×* 3 grid plus one pilot run (PILOT_A1_g5000_c2000_s1), which is scored in this fidelity check and nowhere else in the paper. The 0.10 tolerance in A is a display boundary fixed before the analysis and was not recalibrated in this study. Values for every panel are given in Additional file 2.

### Additional file 1: Fig. S3

**Fig. S3 | The full parameter grid.** (A) OP *R*^2^ for the complete 36-simulation grid (2k, 5k and 10k genes by 0.5k, 2k and 8k cells), one facet per construction; the centre cell holds four simulations and shows their mean. The other eight cells per construction hold one each. (B) Zero-out-degree rate on the same grid and layout. (C) Median top-20 overlap on the same grid. (D) Median number of genes with *P <* 0.05 on the same grid. (E) OP *R*^2^ by number of genes simulated (boxes are median and interquartile range, points are simulations). (F) OP *R*^2^ by number of cells simulated. (G) Number of genes with *P <* 0.05 by construction. (H) OP *R*^2^ against zero-out-degree rate across the 36 simulations; the dashed line is a least-squares fit. (I) Top-20 overlap against zero-out-degree rate, with a least-squares fit. OP *R*^2^ is the out-of-sample prediction *R*^2^ of the knock-out model. Statistics for every panel (*n*, test and exact *P*) are given in Additional file 2.

### Additional file 1: Fig. S4

**Fig. S4 | Anatomy of the zero-out-degree genes.** (A) Distribution of log out-degree for genes with non-zero (pale blue) and zero (pink) out-degree (overlapping histograms, 40 bins); zero-out-degree genes sit at the floor of the scale. (B) log out-degree against log mean expression for all genes; the zero group forms a horizontal band and the dashed line marks the floor. (C) Zero-out-degree rate by gene family (lollipop, gene counts in brackets). (D) Zero-out-degree rate by expression decile, low to high. (E) Zero-out-degree rate by detection-rate decile. (F) Mean shift *Z* after knock-out, zero-out-degree versus non-zero knock-outs (boxes are median and interquartile range, points are genes). (G) Number of zero-out-degree genes against network size for the nine datasets, coloured by species; the dashed line is a least-squares fit. (H) OP *R*^2^ against zero-out-degree rate across the nine datasets; the dashed line is a least-squares fit. (I) Zero-out-degree rate of each dataset (lollipop). The correlation in H is taken across datasets, not within a dataset. Statistics for every panel (*n*, test and exact *P*) are given in Additional file 2.

### Additional file 1: Fig. S5

**Fig. S5 | What it costs.** (A) Network construction time and total knock-out time for the four full-scale runs (two datasets at 2,000 and 8,000 cells per network), joined per run. (B) Setup plus all knock-outs for the same four runs. (C) Wall-clock time against time per knock-out for the six third-party runs, coloured by highly variable gene setting, labelled by GSE number. (D) Reading the network out-degree directly versus knocking out 60 genes one by one (log scale). (E) Simulator generation time against the number of genes simulated (36 simulations, coloured by gene tier). (F) Simulator setup time against time per knock-out for the 36 simulations, coloured by construction. (G) Time per knock-out against zero-out-degree rate across the nine datasets. (H) Time per knock-out against network size across the nine datasets; the dashed line is a least-squares fit. (I) Projected time of a 40-gene screen, 300 s of setup plus 40 times the measured per-knock-out time, for each dataset. Statistics for every panel (*n*, test and exact *P*) are given in Additional file 2.

### Additional file 1: Fig. S6

**Fig. S6 | Data and code resources.** (A) OP *R*^2^ of all 17 tool runs (nine public datasets, five highly variable gene variants and three third-party datasets), ordered by value and coloured by source; the dashed line is 0.80. (B) Metadata matrix for the nine public datasets: knock-outs performed, genes in the network, zero-out-degree rate and OP *R*^2^. Values are printed in the cells and the fill is normalised within each column, so it ranks datasets within a variable and does not compare across variables. (C) Conceptual pipeline, marked conceptual: expression matrix, quality control and ten PCNets by tensor decomposition, then three diagnostics, grade and report. (D) Coverage of the study in log scale: public datasets, third-party datasets, simulated datasets, simulated knock-outs and DepMap knock-outs. (E) Species composition of the nine public datasets, one tile per dataset. (F) Genes retained after quality control (retained over total) for eight datasets; GSE172410 is omitted because its total gene count is not registered in the source table. (G) Knock-out count against OP *R*^2^ for the 17 runs, with a least-squares fit. (H) Knock-out count against network size for the 17 runs. (I) Paired OP *R*^2^ at 5k and 10k highly variable genes for the four datasets that have both settings. The 0.80 line in A is the grade boundary fixed before the analysis and was not recalibrated in this study. Statistics for every panel are given in Additional file 2.

## Notes

### Competing Interest Statement

The authors have declared no competing interest.

