## Supplementary Figures S1-S6 for "Zero-shot virtual knock-out rankings do not recover causal genes from observational single-cell data"

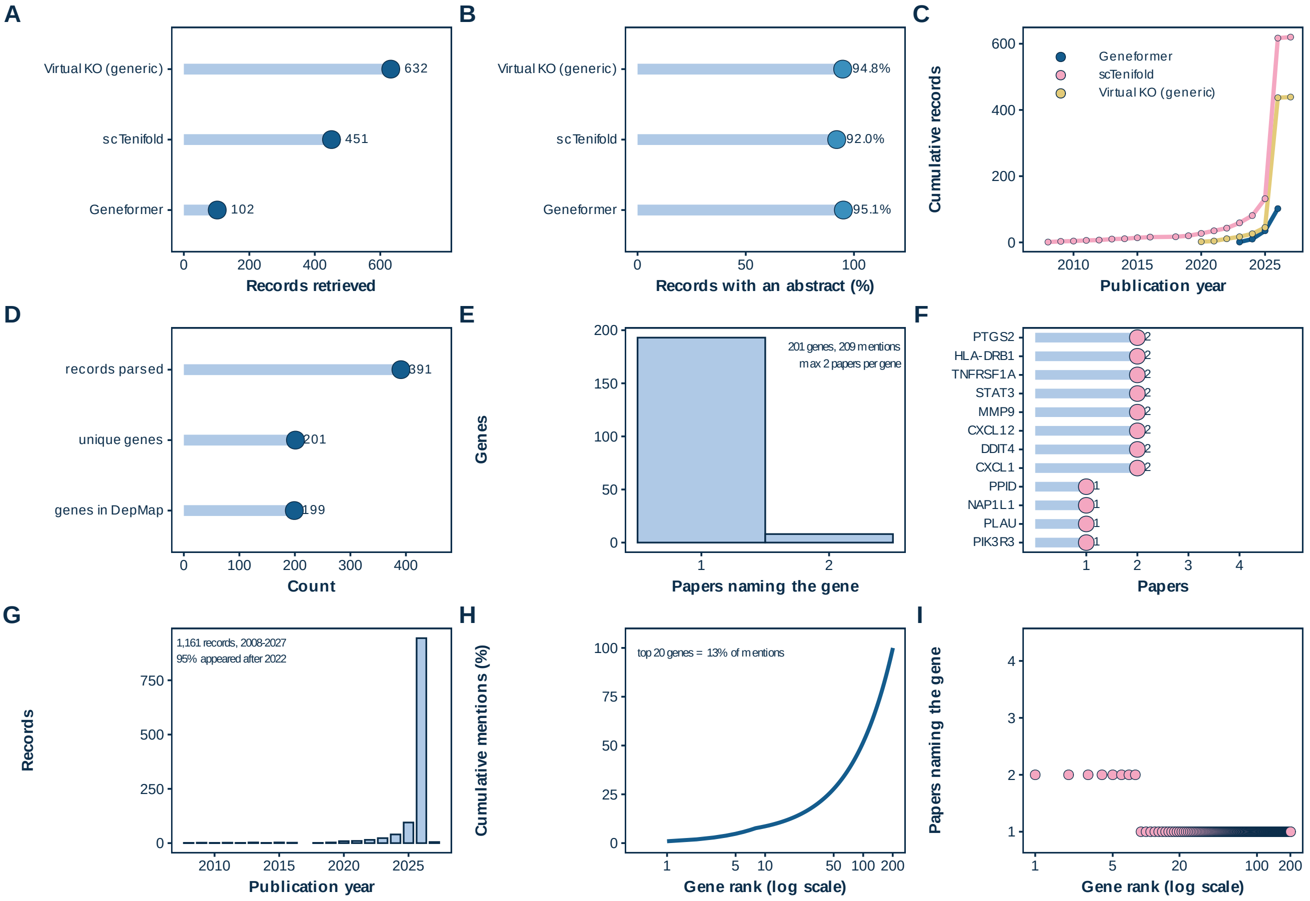

A
Virtual KO (generic)
scTenifold
Geneformer
632
451
102
0
200
400
600
Records retrieved
B
Virtual KO (generic)
scTenifold
Geneformer
94.8%
92.0%
95.1%
0
50
100
Records with an abstract (%)
C
Cumulative records
Geneformer
scTenifold
Virtual KO (generic)
0
200
400
600
2010
2015
2020
2025
Publication year
D
records parsed
unique genes
genes in DepMap
391
201
199
0
100
200
300
400
Count
E
Genes
201 genes, 209 mentions
max 2 papers per gene
0
50
100
150
200
1
2
Papers naming the gene
F
PTGS2
HLA-DRB1
TNFRSF1A
STAT3
MMP9
CXCL12
DDIT4
CXCL1
PPID
NAP1L1
PLAU
PIK3R3
2
1
1
2
3
4
Papers
G
Records
1,161 records, 2008-2027
95% appeared after 2022
0
250
500
750
2010
2015
2020
2025
Publication year
H
Cumulative mentions (%)
top 20 genes = 13% of mentions
0
25
50
75
100
1
5
10
50
100
200
Gene rank (log scale)
I
Papers naming the gene
1
2
3
4
1
5
20
100
200
Gene rank (log scale)

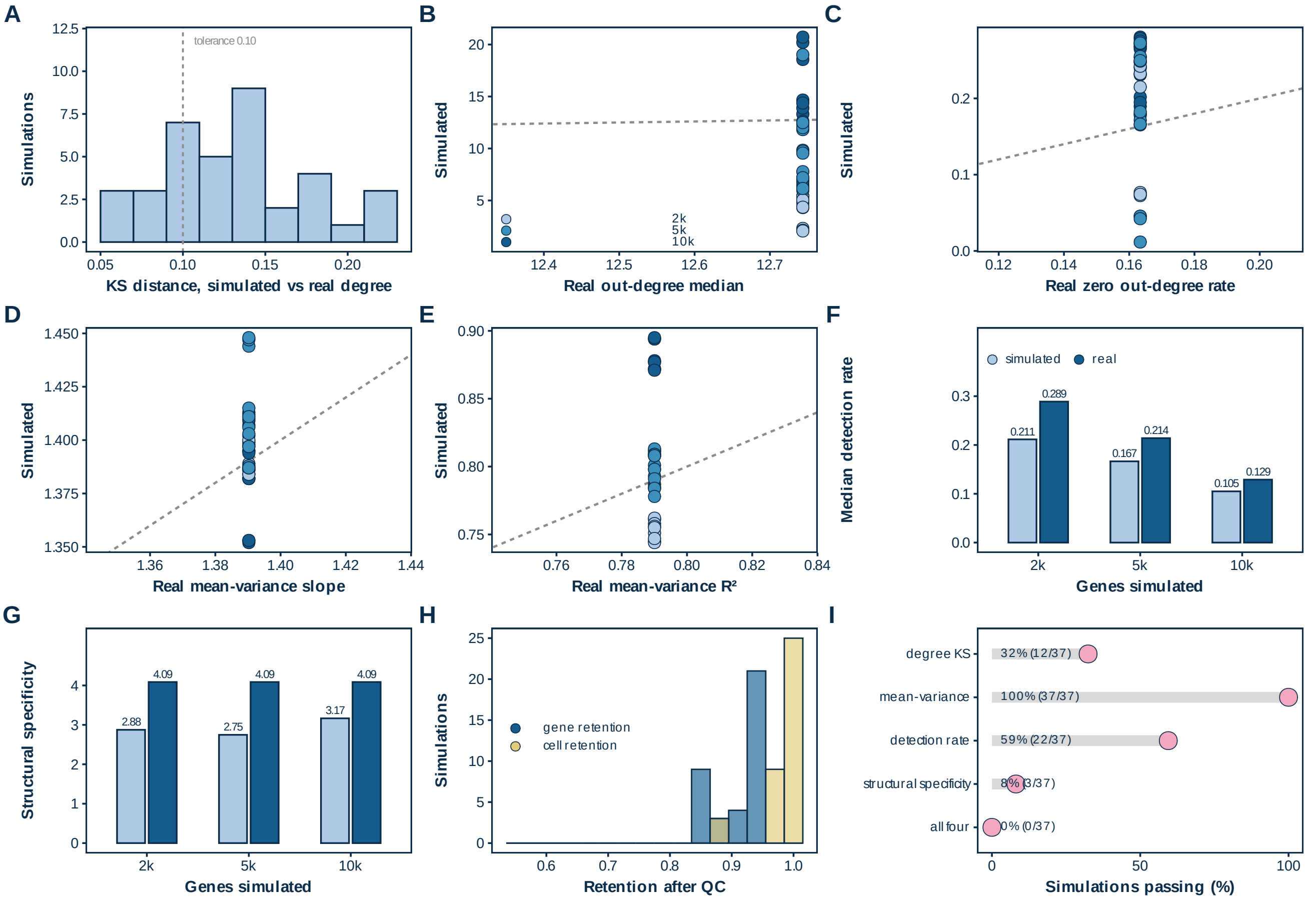

A
tolerance 0.10
Simulations
KS distance, simulated vs real degree
B
Simulated
2k
5k
10k
Real out-degree median
C
Simulated
Real zero out-degree rate
D
Simulated
Real mean-variance slope
E
Simulated
Real mean-variance R²
F
simulated
real
0.211
0.289
0.167
0.214
0.105
0.129
Median detection rate
Genes simulated
G
2.88
4.09
2.75
4.09
3.17
4.09
Structural specificity
Genes simulated
H
gene retention
cell retention
Simulations
Retention after QC
I
degree KS
32% (12/37)
mean-variance
100% (37/37)
detection rate
59% (22/37)
structural specificity
8% (3/37)
all four
0% (0/37)
Simulations passing (%)

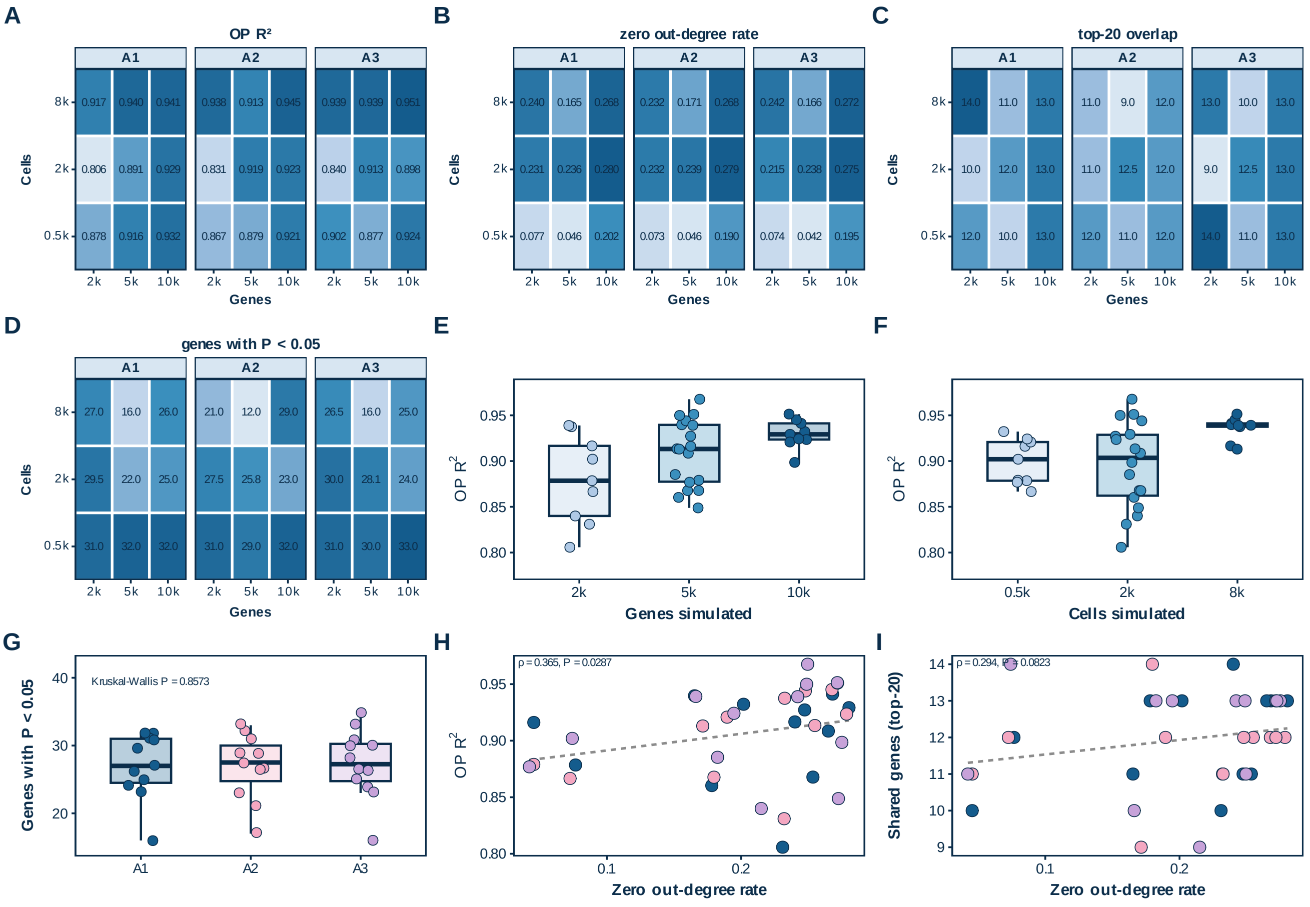
A
OP R²
A1
A2
A3
Cells
8k
2k
0.5k
0.917 0.940 0.941
0.806 0.891 0.929
0.878 0.916 0.932
0.938 0.913 0.945
0.831 0.919 0.923
0.867 0.879 0.921
0.939 0.939 0.951
0.840 0.913 0.898
0.902 0.877 0.924
2k 5k 10k
Genes
B
zero out-degree rate
0.240 0.165 0.268
0.231 0.236 0.280
0.077 0.046 0.202
0.232 0.171 0.268
0.232 0.239 0.279
0.073 0.046 0.190
0.242 0.166 0.272
0.215 0.238 0.275
0.074 0.042 0.195
C
top-20 overlap
14.0 11.0 13.0
10.0 12.0 13.0
12.0 10.0 13.0
11.0 9.0 12.0
11.0 12.5 12.0
12.0 11.0 12.0
13.0 10.0 13.0
9.0 12.5 13.0
14.0 11.0 13.0
D
genes with P < 0.05
27.0 16.0 26.0
29.5 22.0 25.0
31.0 32.0 32.0
21.0 12.0 29.0
27.5 25.8 23.0
31.0 29.0 32.0
26.5 16.0 25.0
30.0 28.1 24.0
31.0 30.0 33.0
E
OP R²
Genes simulated
F
Cells simulated
G
Genes with P < 0.05
Kruskal-Wallis P = 0.8573
H
ρ = 0.365, P = 0.0287
Zero out-degree rate
I
ρ = 0.294, P = 0.0823
Shared genes (top-20)
Zero out-degree rate

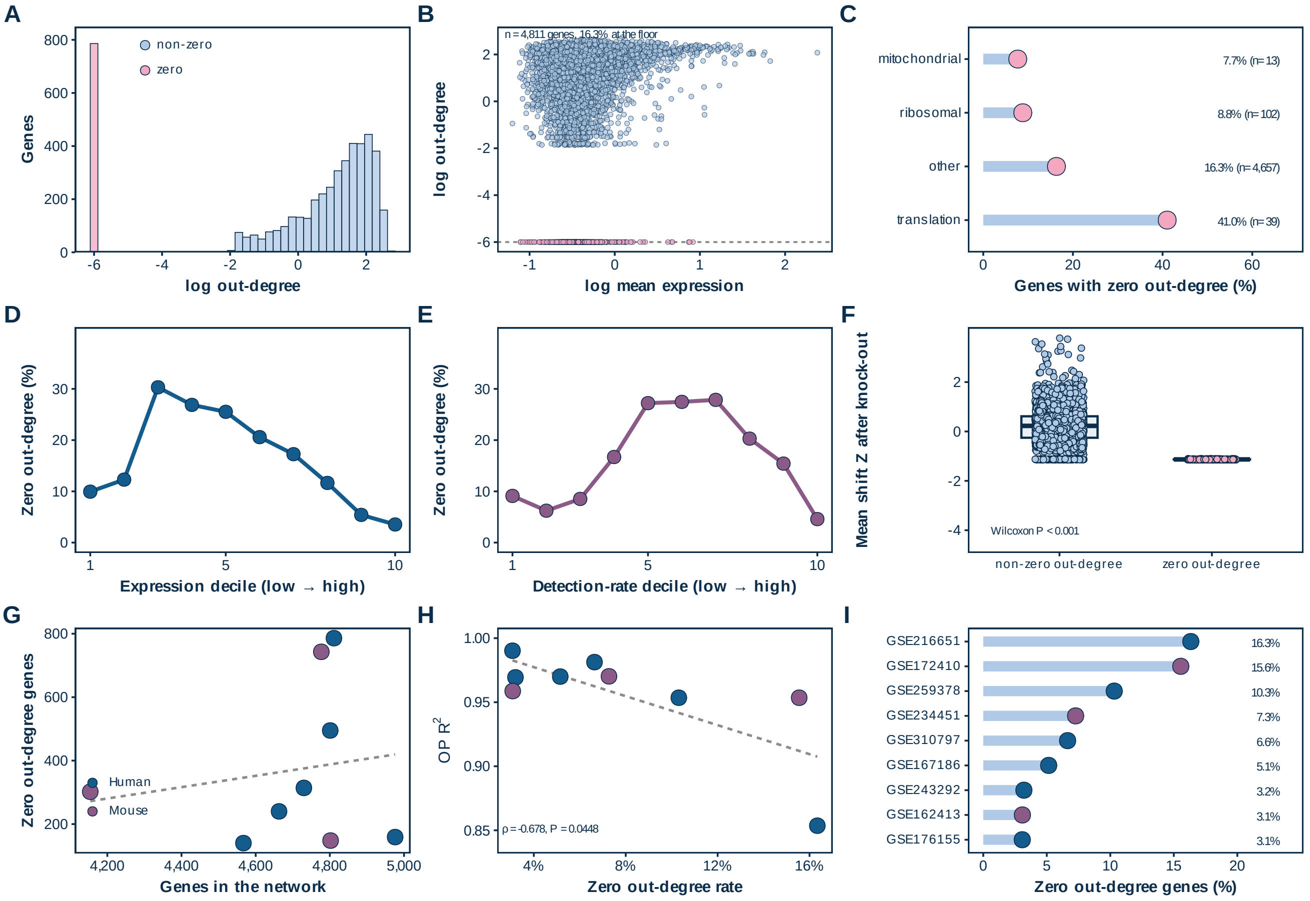

A
Genes
log out-degree
non-zero
zero
B
n = 4,811 genes, 16.3% at the floor
log out-degree
log mean expression
C
mitochondrial
ribosomal
other
translation
7.7% (n = 13)
8.8% (n = 102)
16.3% (n = 4,657)
41.0% (n = 39)
Genes with zero out-degree (%)
D
Zero out-degree (%)
Expression decile (low → high)
E
Zero out-degree (%)
Detection-rate decile (low → high)
F
Mean shift Z after knock-out
Wilcoxon P < 0.001
non-zero out-degree
zero out-degree
G
Zero out-degree genes
Human
Mouse
Genes in the network
H
OP R2
ρ = -0.678, P = 0.0448
Zero out-degree rate
I
GSE216651
GSE172410
GSE259378
GSE234451
GSE310797
GSE167186
GSE243292
GSE162413
GSE176155
16.3%
15.6%
10.3%
7.3%
6.6%
5.1%
3.2%
3.1%
3.1%
Zero out-degree genes (%)

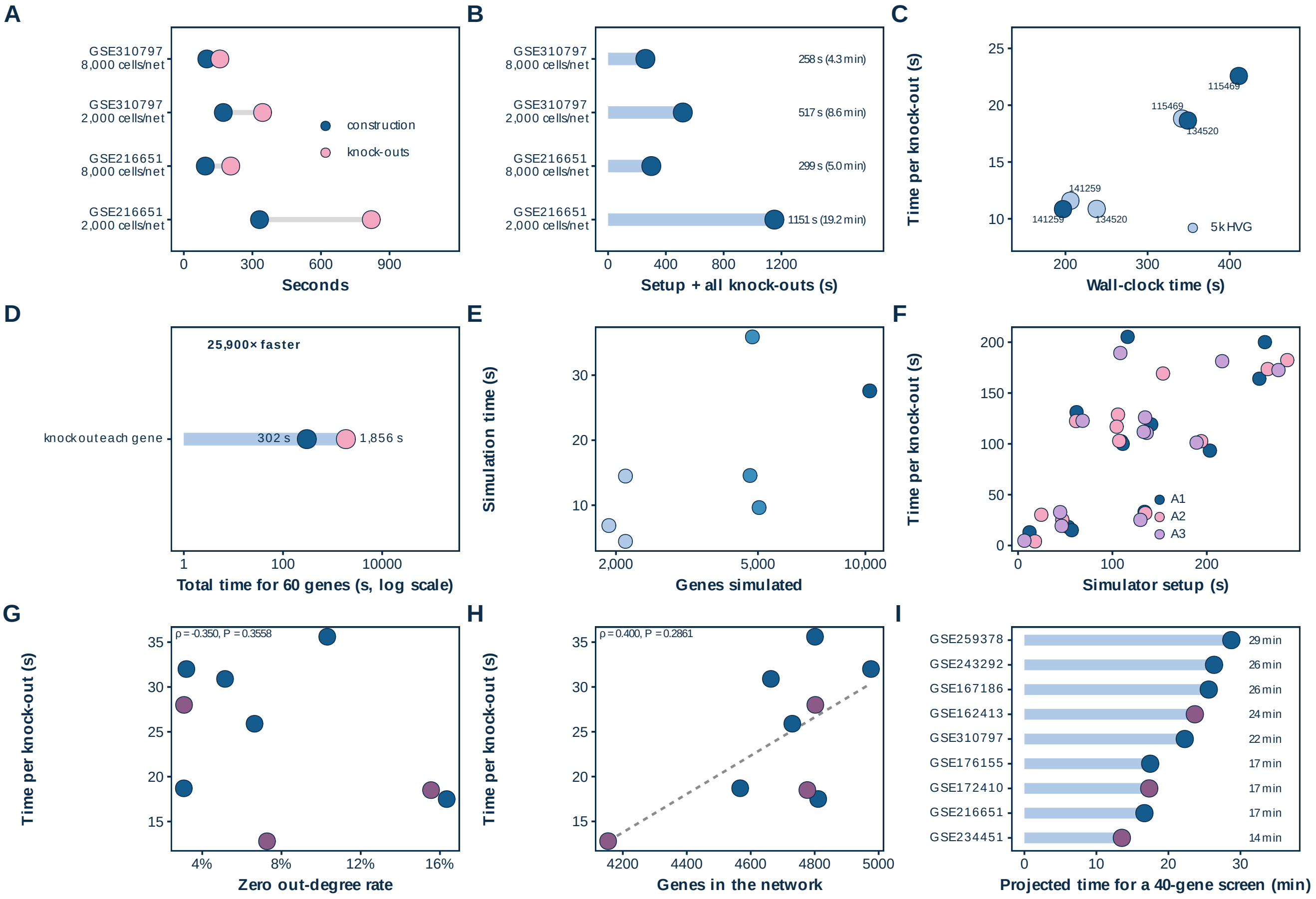
A
GSE310797
8,000 cells/net
GSE310797
2,000 cells/net
GSE216651
8,000 cells/net
GSE216651
2,000 cells/net
construction
knock-outs
0
300
600
900
Seconds
B
258 s (4.3 min)
517 s (8.6 min)
299 s (5.0 min)
1151 s (19.2 min)
0
400
800
1200
Setup + all knock-outs (s)
C
Time per knock-out (s)
115469
134520
141259
5k HVG
Wall-clock time (s)
D
25,900× faster
knockout each gene
302 s
1,856 s
Total time for 60 genes (s, log scale)
E
Simulation time (s)
2,000
5,000
10,000
Genes simulated
F
A1
A2
A3
Simulator setup (s)
G
ρ = -0.350, P = 0.3558
Zero out-degree rate
H
ρ = 0.400, P = 0.2861
Genes in the network
I
GSE259378
GSE243292
GSE167186
GSE162413
GSE310797
GSE176155
GSE172410
GSE216651
GSE234451
29 min
26 min
26 min
24 min
22 min
17 min
17 min
17 min
14 min
Projected time for a 40-gene screen (min)

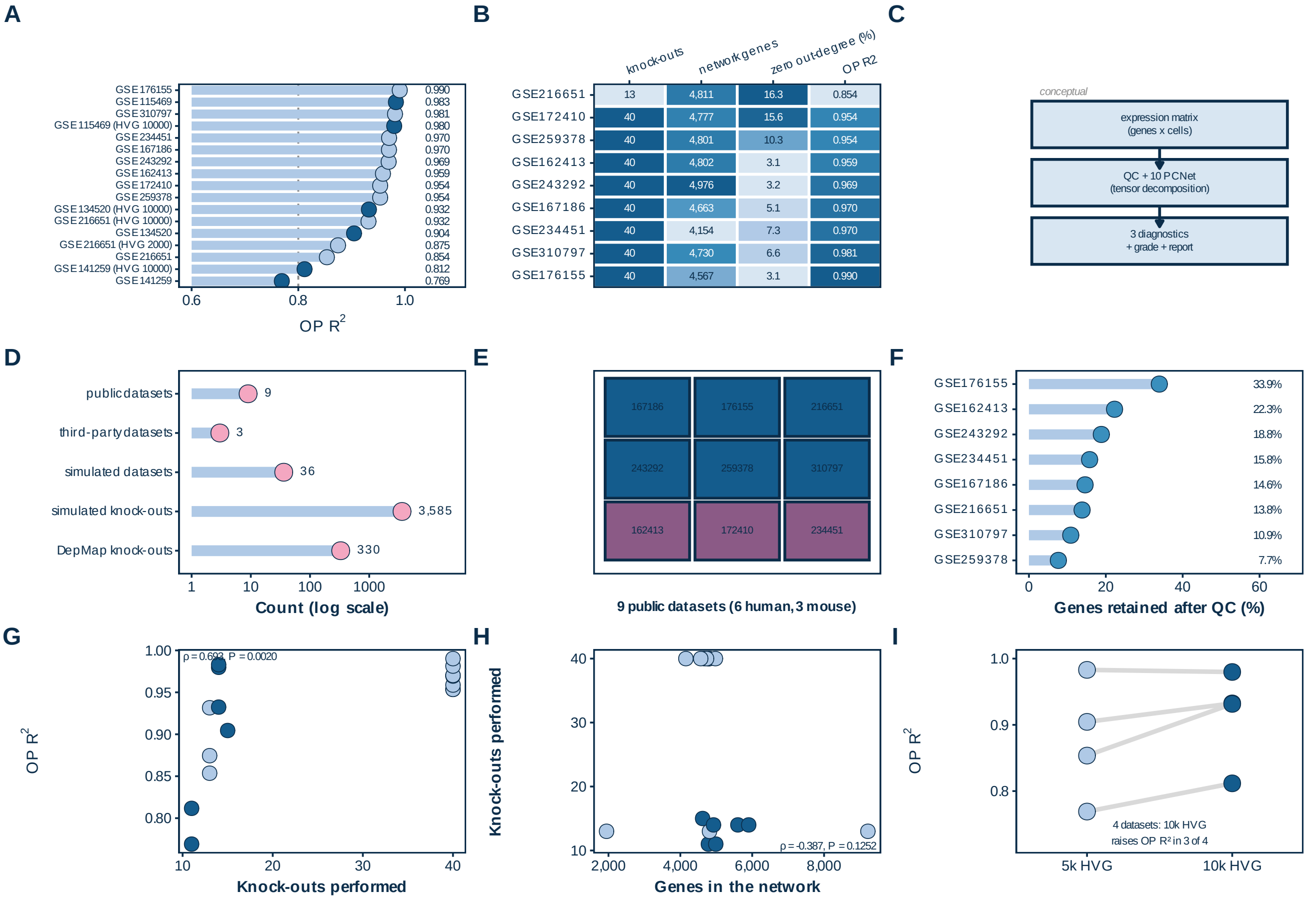
A
GSE176155
GSE115469
GSE310797
GSE115469 (HVG 10000)
GSE234451
GSE167186
GSE243292
GSE162413
GSE172410
GSE259378
GSE134520 (HVG 10000)
GSE216651 (HVG 10000)
GSE134520
GSE216651 (HVG 2000)
GSE216651
GSE141259 (HVG 10000)
GSE141259
0.990
0.983
0.981
0.980
0.970
0.970
0.969
0.959
0.954
0.954
0.932
0.932
0.904
0.875
0.854
0.812
0.769
0.6
0.8
1.0
OP R2
B
knock-outs
network genes
zero out-degree (%)
OP R2
GSE216651 13 4,811 16.3 0.854
GSE172410 40 4,777 15.6 0.954
GSE259378 40 4,801 10.3 0.954
GSE162413 40 4,802 3.1 0.959
GSE243292 40 4,976 3.2 0.969
GSE167186 40 4,663 5.1 0.970
GSE234451 40 4,154 7.3 0.970
GSE310797 40 4,730 6.6 0.981
GSE176155 40 4,567 3.1 0.990
C
conceptual
expression matrix
(genes x cells)
QC + 10 PCNet
(tensor decomposition)
3 diagnostics
+ grade + report
D
public datasets
third-party datasets
simulated datasets
simulated knock-outs
DepMap knock-outs
9
3
36
3,585
330
1
10
100
1000
Count (log scale)
E
167186
176155
216651
243292
259378
310797
162413
172410
234451
9 public datasets (6 human, 3 mouse)
F
GSE176155
GSE162413
GSE243292
GSE234451
GSE167186
GSE216651
GSE310797
GSE259378
33.9%
22.3%
18.8%
15.8%
14.6%
13.8%
10.9%
7.7%
0
20
40
60
Genes retained after QC (%)
G
ρ = 0.693, P = 0.0020
OP R2
1.00
0.95
0.90
0.85
0.80
10
20
30
40
Knock-outs performed
H
Knock-outs performed
40
30
20
10
2,000
4,000
6,000
8,000
ρ = -0.387, P = 0.1252
Genes in the network
I
OP R2
1.0
0.9
0.8
4 datasets: 10k HVG
raises OP R² in 3 of 4
5k HVG
10k HVG
